# Hormetic elevation of taurine restrains inflammaging by deactivating the NLRP3 inflammasome

**DOI:** 10.1101/2025.05.27.656381

**Authors:** Chenyang Guan, Seungjin Ryu, Mingze Dong, Yun-Hee Youm, Subhasis Mohanty, Rae Maeda, Lucie Orliaguet, Hee-Hoon Kim, Tamara Dlugos, Steven R. Smith, Eric Ravussin, Janset Onyuru, Andrew Wang, Albert C. Shaw, Hal M. Hoffman, Yuval Kluger, Yuki Sugiura, Vishwa Deep Dixit

## Abstract

Taurine, the most abundant sulfonic amino acid in humans is largely obtained from diets rich in animal proteins. However, taurine is dietary non-essential because it can be synthesized from cysteine by activation of transsulfuration pathway (TSP) when food consumption is low or if the diet is predominantly plant based. The decline of taurine was proposed as the driver of aging through an undefined mechanism. Here, we found that mild food restriction in humans for one year that resulted in 14% reduction of calorie intake elevated the hypotaurine and taurine concentration in adipose tissue. Therefore, we investigated whether elevated taurine mimics caloric-restriction’s beneficial effects on inflammation, a key mechanism of aging. Interestingly, aging increased the circulating and tissue concentrations of taurine suggesting that elevated taurine may serve as a hormetic stress response metabolite that regulates mechanism of age-related inflammation. The elevated taurine protected mice against mortality from sepsis and inhibited inflammasome-driven inflammation and gasdermin-D (GSDMD) mediated pyroptosis. Mechanistically, ‘danger signals’ including hypotonicity that activate NLRP3-inflammasome, caused upstream taurine efflux from macrophages, which triggered potassium (K^+^) release and downstream canonical NLRP3 inflammasome assembly, caspase-1 activation, GSDMD cleavage and IL-1β and IL-18 secretion that was reversed by taurine restoration. Notably, taurine does not efflux from GSDMD pore and inhibited IL-1β from macrophages independently of known transporters SLC6A6 and SLC36A1. Increased taurine in old mice promotes healthspan by inducing anti-inflammatory pathways previously linked to youthfulness. These findings demonstrate that taurine is an upstream metabolic sensor of cellular perturbations that control NLRP3 inflammasome and lowers age-related inflammation.

## Main

Aging related chronic diseases are linked with increased inflammation that is driven in part by NLRP3 inflammasome activation ^1–3^. The inflammasome-dependent elevation of interleukin (IL)-1β and IL-18 acts on multiple cell types ^4^ , including but not limited to neurons ^5^, stem cells ^6^ and adipocytes ^7–9^ leading to functional decline in aging also described as “inflammaging” ^1–3,10–12^. Mangum and Towle proposed a physiological adaptation state termed “enantiostasis”, where despite an unstable internal cellular milieu, the net effect of regulatory flux is stability or tendency towards it ^13^. Thus, to stall precipitous functional decline during aging, enantiostatic endogenous molecules may need to be mobilized to oppose mechanisms that cause instability. Immunometabolic reprogramming induced by calorie restriction(CR)-induced negative balance in humans upregulates protective pathways that restrains inflammaging ^14^ and may control longevity^15^. The identity of CR-induced enantiostatic metabolites that inhibit mechanism of age-related inflammation remains unknown.

Taurine, the most abundant non synthetic zwitterionic sulfonic amino acid is largely obtained from diets rich in animal proteins with intracellular levels in cells ranging from 5-50mM ^16,17^. However, taurine is dietary non-essential because it can be synthesized from cysteine if diet is taurine deficient ^18^. The restriction of sulfur amino acids that activates the dormant transsulfuration pathway to maintain endogenous cysteine and taurine is linked to longevity in mice ^19,20^. Decline in taurine was recently proposed as the driver of aging process ^21^. Multiple longevity interventions in rodents are associated with upregulation of the transsulfuration pathway (TSP) enzyme cystathionine γ-lyase (CTH) ^22^, which converts cystathionine into cysteine ^23^. Cysteine is also a key precursor for synthesis of taurine; the enzyme cysteine dioxygenase (CDO) converts cysteine into cysteine sulfinate, which is further decarboxylated into hypotaurine by cysteine sulfinate decarboxylase (CSAD). The enzymes of the flavin monooxygenase (FMO) family converts hypotaurine to generate taurine ^24–26^. It is thought that mammals are unable to oxidize the sulfur in taurine or cleave its C-S bond. Thus, taurine is thought to be a final product of cysteine metabolism downstream of TSP and is largely excreted in its biochemically native state or as conjugates with bile acids or xenobiotics for elimination ^16^. Given its zwitterionic nature, taurine is highly water soluble and its concentration in leukocytes is reported to be as high as 20 mM ^27,28^. In addition, taurine is implicated in dampening inflammation and maintenance of cell volume by serving as organic osmolyte ^29^, however the unifying mechanism of taurine’s action on innate immune sensing pathways regulating inflammaging are unknown.

## Caloric restriction in humans rewires transsulfuration pathway to elevate taurine

The <u>C</u>omprehensive <u>A</u>ssessment of <u>L</u>ong-term <u>E</u>ffects of <u>R</u>educing Intake of <u>E</u>nergy (CALERIE-II) clinical trial in healthy adults demonstrated that food restriction leading to 14% reduction in caloric intake lowers inflammation ^14,30^. To determine the metabolites that may function as signaling regulators of CR’s anti-inflammatory response, we conducted unbiased metabolomics analyses of subcutaneous adipose tissue (SAT) of participants in the CALERIE-II trial at baseline and one year after CR (Fig. 1a). The top 30 up-and down-regulated metabolites in human adipose tissue revealed that CR in humans is associated with a reduction in cysteine and GSH, but a surprising increase in hypotaurine and taurine, the terminal products of cysteine metabolism in TSP (Fig. 1a). Further pathway analyses revealed that taurine and hypotaurine metabolism were the core pathway upregulated upon 1 year of CR in humans (Fig. 1b, c and Fig. S1a). There was also a trend toward an increase in cysteine sulfinic acid (p = 0.0768), which is synthesized from cysteine by the action of CDO, with no significant change in homocysteine, cystathionine, bisulfite or pyruvate (Fig. S1b, c). Consistent with these data, the RNA sequencing analyses of SAT found increased expression of *Csad* and *Fmo1*, the enzymes responsible for generating hypotaurine and taurine (Fig. 1d). Also, similar to *Fmo1*, CR significantly increased the expression of *Fmo2* at year 1 (Fig. 1d), which is previously shown to extend lifespan in *Caenorhabditis elegans* ^31,32^. There was no change in the expression of *Cdo1* or major taurine transporter *Slc6a6* (Fig. S1d). CR significantly increased the concentration of cystathionine and lowered the redox regulator GSH in blood (Fig. S1e). Notably, the elevation of taurine and hypotaurine were specific to the human adipose tissue as CR did not affect these metabolites in the plasma (Fig. S1f). The CR-induced increase in taurine in adipose tissue was also linked to significant decrease in inflammasome pathway genes *Il-18, Asc, Nek7* with a trend in reduction of *Nlrp3* (Fig. S1g). Together, these findings demonstrate that chronic mild food restriction in humans reduces cysteine and its product GSH, with an increase in taurine concentration in adipose tissue. These findings suggest that CR-induced activation of transsulfuration pathway (TSP) is associated with lower inflammation and prioritization of metabolic demand that favors synthesis of hypotaurine and taurine downstream of cysteine.

**Fig. 1.**
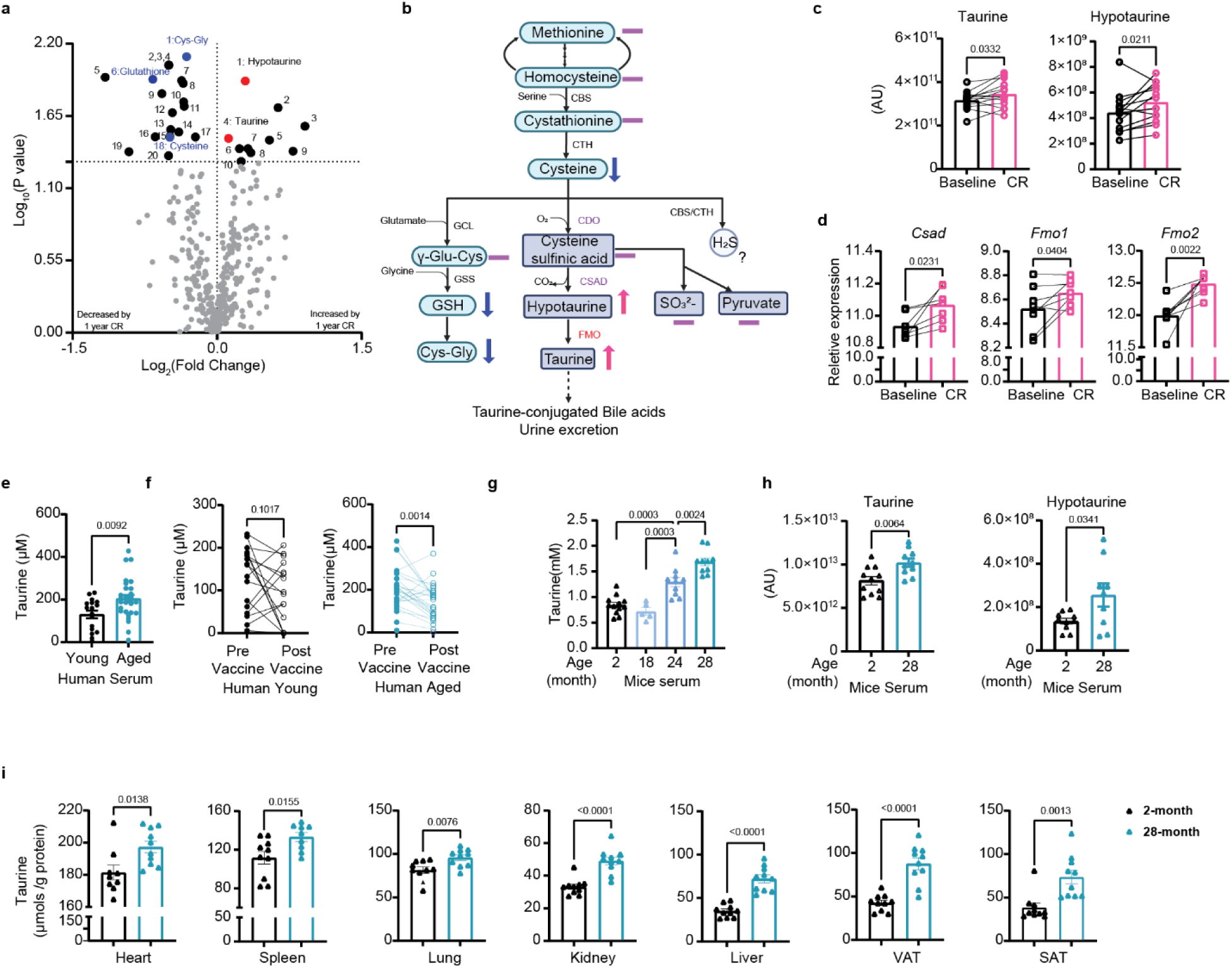
Calorie restriction and aging elevate taurine levels. **(a-c)** Unbiased metabolomics analyses of subcutaneous adipose tissue (SAT) of participants in the CALERIA-II trail at baseline and one year after caloric restriction (CR). (a) Volcano plots of the 387 metabolites of SAT from healthy individuals at baseline and 1-year CR. Each dot represents a metabolite identified by LC-MS/MS. The volcano plot shows the fold-change (x-axis) versus the significance (y-axis) of the identified 387 metabolites. The significance (non-adjusted P-value) and the fold change are converted to -Log_10_(P-value) and log_2_(Fold-change). The vertical and horizontal dotted lines show the cut-off of fold-change= ±1.0, and of p-value=0.05. 30 significantly regulated metabolites at 1 year of CR compared with baseline were heighted (up-regulated in red, down-regulated in blue (n=14. P<0.05). (b) Schematic summary of the transsulfuration pathway and metabolites from baseline to 1 year CR that were measured in the healthy human SAT. Blue arrows indicate significantly decreasing, pink arrows indicate significantly increasing metabolites and purple lines indicate not significant changes. (c) Changes in the levels of taurine and hypotaurine measured by LC-MS/MS in human SAT after 1 year CR. (Significance was calculated using paired t-test). **(d)** The RNA-sequencing analyses of SAT of participants in the CALERIA-II trail at baseline and after 1 year CR. The expression level of *Csad, Fmo1* and *Fmo2* after CR. Paired t-test was performed (n=8). **(e)** Circulating taurine concentration from serum in young and aged humans (n=17, 28). **(f)** Taurine level changes prior and 2 days after influenza vaccine immunization in young and aged human serum (young n=17, aged n=28). **(g)** Serum taurine levels in different ages of male C57BL/6J mice (n=12, 5, 10, 12, Yale rodent colony), mice are littermates. **(h)** Serum taurine and hypotaurine levels were determined in 2-month-old and 28-month-old male C57BL/6N mice (n=11/10) by LC-MS/MS. (**i)** Tissues taurine levels in 2-month-old and 28-month-old C57BL/6J male mice (n=10/group), mice are littermates, by taurine assay kit. Error bars represent the mean ± SEM.

## Circulating and tissue concentration of taurine increase and doesn’t decline with aging

Since CR increases lifespan and elevates taurine levels, one can posit that aging maybe associated with a decline in taurine in mice and humans. However, in contrast to a recent study that reported reduction of taurine in older adults ^21^, our cross-sectional analyses found that in humans, aging is associated with a significant elevation in taurine concentrations (Fig. 1e). At 2 days after vaccination, the young study participants demonstrated no change while older adults displayed a significant reduction in taurine concentrations (Fig. 1f). Consistent with human data, aged C57BL/6 mice (Yale animal colony) also displayed higher taurine concentrations in serum (Fig. 1g). This was orthogonally validated using metabolomics analyses of sera by LC/MS in a separate cohort of mice (NIA rodent colony) which further confirmed that aging is associated with higher taurine and hypotaurine (Fig. 1h), but lower homocysteine, cystathionine, cysteine sulfinic acid with no changes of cysteine concentration in wild type mice (Fig. S1h). Interestingly, taurine concentration is also high in multiple tissues (Fig. S1i). We found that similar to serum, aging was associated with significant elevation of taurine concentrations in heart, spleen, lung, kidney, liver, visceral adipose tissue (VAT) and subcutaneous adipose tissue (SAT), but not in brown adipose tissue (BAT) and muscle (Fig. 1i, Fig. S1j). These results suggest that aging is not caused by taurine deficiency and that increases of taurine observed post-CR in healthy adults and in older mice and humans could be adaptive hormetic-stress response designed to restore homeostasis.

## Upstream taurine efflux in response to danger signals activates the NLRP3 inflammasome

Given the high mM cellular and tissue concentration of taurine^28^ and a critical role of macrophages in control of inflammaging, we tested whether taurine controls central mechanisms of inflammation such as the NLRP3 inflammasome. Sensing of multiple ‘danger signals’ such as extracellular ATP ^33^, urate crystals ^34^, ceramides ^35^, oxidized cardiolipin and ectopic DNA from mitochondria ^36,37^ can activate the NLRP3 inflammasome which cleaves caspase-1 to release IL-1β and IL-18 ^38^. The unbiased LC/MS metabolomics (Fig. S2a, 2b) revealed that NLRP3 inflammasome activation in macrophages increases glycolytic metabolites ^39^ catalyzed by phosphofructokinase-1 ^40^ , such as fructose 1, 6-biphosphate together with significant reduction in nicotinamide adenine dinucleotide (NAD) (Fig. 2a), a key cofactor in redox reactions^41^ with no change in cystine (oxidized form of cysteine) and betaine an organic osmolyte in methionine cycle (Fig. 2b, S2c). Interestingly, we found that NLRP3 inflammasome activation caused significant reduction in the concentration of taurine (Fig. 2b) with marked changes in pathways regulating pentose phosphate pathway, arginine and proline and citrulline metabolism (Fig 2a, Fig S2b). Notably, reduction of taurine in NLRP3 inflammasome activated macrophages did not alter GSH and oxidized GSH, suggesting no change in glutathione redox potential (E_GSH_) (Fig. S2d). The intracellular taurine concentration in murine bone marrow derived macrophages (BMDMs) is approximately 10mM and comparable to taurine rich organs such as heart and muscle (Fig. S2e)^27^. Interestingly, multiple activators of NLRP3 such as ATP, hypotonicity, monosodium urate crystals (MSU), ceramides, nigericin and imiquimod caused varying degree of taurine efflux (Fig. 2c, Fig. S2f). Further quantification of taurine levels confirmed that extracellular ATP caused rapid and significant efflux of intracellular taurine from macrophages with accumulation in the cell supernatants (Fig. 2d).

**Fig. 2.**
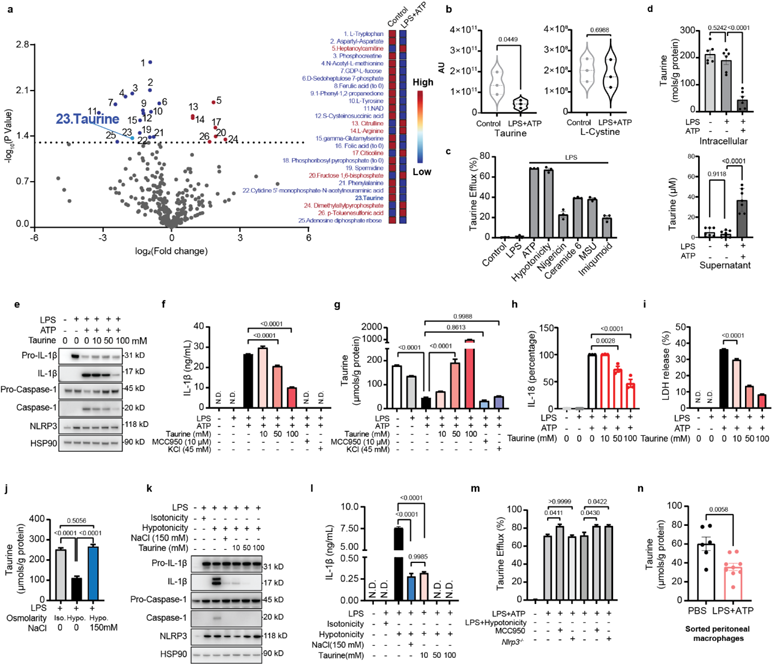
Taurine inhibits NLRP3 inflammasome activation. **(a)** Volcano plots of the 290 metabolites identified between the LPS plus ATP challenge group and control group in the bone marrow derived macrophages (BMDMs). Each dot represents a metabolite identified by LC-MS/MS. The volcano plot shows the fold-change (x-axis) versus the significance (y-axis) of the identified 290 metabolites. The significance (non-adjusted p-value) and the fold change are converted to -Log_10_(p-value) and log_2_(fold-change). The horizontal dotted lines show the cut-off of fold-change= ±1.0, and of p-value=0.05. 26 significantly regulated metabolites in response to LPS plus ATP treatment compared with baseline were heighted (up-regulated in red, down-regulated in blue (n=3, P<0.05). The right panel is the top 26 discriminating parameters in descending order of importance from the metabolomics data of BMDMs with LPS, ATP treatment and control group. The colored legend on the right indicates the relative abundance of variables, with red and blue indicating high and low values, respectively, while beige illustrates neutral values. **(b)** Changes in the levels of taurine and L-cystine in response to LPS and ATP stimulation in BMDMs, measured by LC-MS/MS. **(c)** Taurine efflux was determined in LPS-primed-BMDMs treated with 10 mM ATP, Hypotonic solution (90 mOsmolarity), 10 μM Nigericin for 45 mins, Ceramide 6, MSU for 6 hours and Imiquimod for 2 hours. **(d)** Intracellular and culture medium taurine levels in BMDMs primed with LPS and stimulated with ATP. **(e)** Western blots of cell lysates and supernatants from BMDMs stimulated with LPS and ATP and treated with taurine. These results are representative of five independent experiments. **(f)** ELISA analysis of IL-1β production from BMDMs stimulated with LPS (1 μg/ml, 4 hours), followed by ATP (5 mM, 45 min) treatment with or without taurine, MCC950 or KCl. **(g)** Intracellular taurine level following inflammasome activation in BMDMs supplemented with 100 mM taurine, 10 μM MCC950 or 45 mM KCl. (Representative of three experiments). **(h-i)** Production of IL-18 and LDH from BMDM stimulated with LPS and ATP and treated with taurine as measured by ELISA (h) and LDH assay (i). **(j)** Intracellular taurine concentration in LPS-primed BMDMs stimulated by hypotonicity in the presence of sodium chloride. (Representative of three experiments). **(k)** Western blots of cell lysates and supernatants from BMDMs activated with hypotonic solution in the presence of taurine or NaCl. (Representative of three experiments). **(l)** Production of IL-1β from BMDMs stimulated with LPS and hypotonic solution and treated with taurine. **(m)** Intracellular taurine level of LPS-primed BMDMs from control and Nlrp3 deficient mice following ATP or hypotonic treatment supplemented with 10 μM MCC950. (Representative of three experiments). **(n)** Intracellular taurine concentration in thioglycolate elicited macrophages sorted from peritoneal cavity of mice treated with LPS and ATP. These data are representative of three independent experiments. Error bars represent the mean ± SEM.

Given an association between taurine efflux and inflammasome activation, we next tested whether elevation of taurine dampens the inflammasome-mediated inflammation in macrophages. Interestingly, restoration of taurine in NLRP3 inflammasome activated macrophages dose-dependently inhibited the IL-1β p17 secretion (Fig. 2e, 2f) and caspase-1 cleavage without affecting the NLRP3 expression (Fig. 2e), with active taurine uptake when taurine was restored (Fig. 2g). Furthermore, the release of IL-18 and lactate dehydrogenase (LDH) induced by NLRP3 activation was blocked by taurine in a dose dependent manner (Fig. 2h, 2i). In addition to extracellular ATP, we confirmed that taurine also inhibited the NLRP3 inflammasome activation in response to monosodium urate crystals (MSU) (Fig. S2g, S2h) and lipotoxic ceramide (Fig. S2i, S2j).

We next determined how taurine efflux impacts inflammasome activation. It is known that upstream efflux of K^+^ from macrophages in response to ‘danger signals’ causes the downstream assembly and activation of NLRP3 inflammasome^42^. Interestingly, taurine efflux in LPS primed and extracellular ATP activated macrophages was not rescued by either restoration of K^+^ or inhibition of NLRP3 by its structural antagonist MCC950^43^ (Fig. 2g), suggesting that taurine efflux is an upstream event that activates downstream NLRP3 inflammasome. As expected KCl, MCC950 and taurine restoration in NLRP3 inflammasome activated cells inhibited IL-1β release (Fig. 2f).

Taurine is also known to function as an organic osmolyte to control cell volume^21^. Notably, the NLRP3 inflammasome can sense alterations in cell osmolarity and resultant changes in cytosolic or membrane integrity ^44,45^. We found that TLR4 primed macrophages exposed to hypotonic stress, a known activator of NLRP3 inflammasome also caused efflux of taurine (Fig. 2j). Interestingly, restoration of isotonicity in media with sodium chloride (NaCl) reversed intracellular taurine concentrations to normal in macrophages (Fig. 2j), suggesting that taurine is a sensor of cytosolic integrity that signals to NLRP3. Consistent with this hypothesis, in LPS primed cells, hypotonicity induced NLRP3 inflammasome activation was inhibited by addition of NaCl that restores isotonicity in media also reverses taurine depletion (Fig. 2j, 2k, 2l). In support of critical role of taurine in defense of cytosolic homeostasis, we found that elevation of taurine in macrophages despite reduced NaCl mediated hypotonic stress, deactivated the NLRP3 mediated caspase-1 cleavage and blocked IL-1β secretion from macrophages (Fig. 2k, 2l).

Consistent with upstream role of taurine in inflammasome regulation, NLRP3 deficiency or its inhibition with small molecule MCC950, did not affect the ATP and hypotonicity-induced taurine efflux from inflammasome primed macrophages (Fig. 2m). We next elicited peritoneal macrophages with thioglycolate treatment in adult mice followed by challenge with LPS and ATP to activate NLRP3 inflammasome. Inflammasome activation in macrophages *in vivo* also led to taurine efflux (Fig. 2n). Taken together, these results demonstrate that taurine serves as an upstream danger sensor and its efflux in response to hypotonicity and DAMPs, links metabolic-innate immune sensing to NLRP3 inflammasome activation in macrophages.

## Taurine deactivates canonical NLRP3 inflammasome and gasdermin-D in macrophages

We further investigated the mechanism and specificity of taurine’s anti-inflammatory action mediated via inflammasome deactivation in macrophages. The NLRP3 inflammasome activation also effluxed folic acid from macrophages (Fig. 2a). However, neither folic acid nor amino acids glycine and glutamine that can impact macrophage function affected the IL-1β secretion in response to NLRP3 inflammasome activation (Fig. S2k-m). Moreover, taurine did not affect tumor necrosis factor–α (TNF-α) secretion from macrophages in response to extracellular ATP (Fig. S3a), MSU (Fig. S3b) or C6 Ceramide (Fig. S3c) suggesting specificity. Taurine also did not affect the inflammasome priming step, as it did not inhibit nuclear factor (NF)-κB activation, Erk, P38 or Jnk signaling in response to toll-like receptor (TLR)-4 activation in macrophages (Fig. S3d).

Next, we examined whether taurine could also inhibit the activation of other inflammasome complexes. The non-NLR AIM2 inflammasome was examined by transfecting BMDMs with the dsDNA analog Poly(dA:dT). The AIM2 inflammasome activation with LPS and Poly(dA:dT) did not cause taurine efflux from macrophages (Fig. 3a). Consistent with this, elevation of taurine did not affect caspase-1 cleavage (Fig. 3b), IL-1β (Fig. 3c) or IL-18 (Fig. S3e) from BMDMs in response to AIM2 inflammasome stimulation. In addition, the gram-negative bacteria derived intracellular LPS is sensed by noncanonical pathway that results in caspase-11–dependent pyroptotic cell death and IL-1β production ^46^. We activated caspase-11 in BMDMs primed with the TLR1/2 ligand Pam3CSK4 and transfected with LPS. The treatment of BMDMs with taurine at the same time with LPS transfection has no inhibitory effect on caspase-1 or IL-1β release (Fig. 3d, Fig. S3f) and did not cause taurine efflux (Fig. 3e). In addition, in LPS primed macrophages bacterial flagellin which activates NLRC4 inflammasome, does not affect taurine efflux or IL-1β (p17) (Fig. S3g). These results indicate selectivity of taurine for regulating NLRP3 inflammasome.

**Fig. 3.**
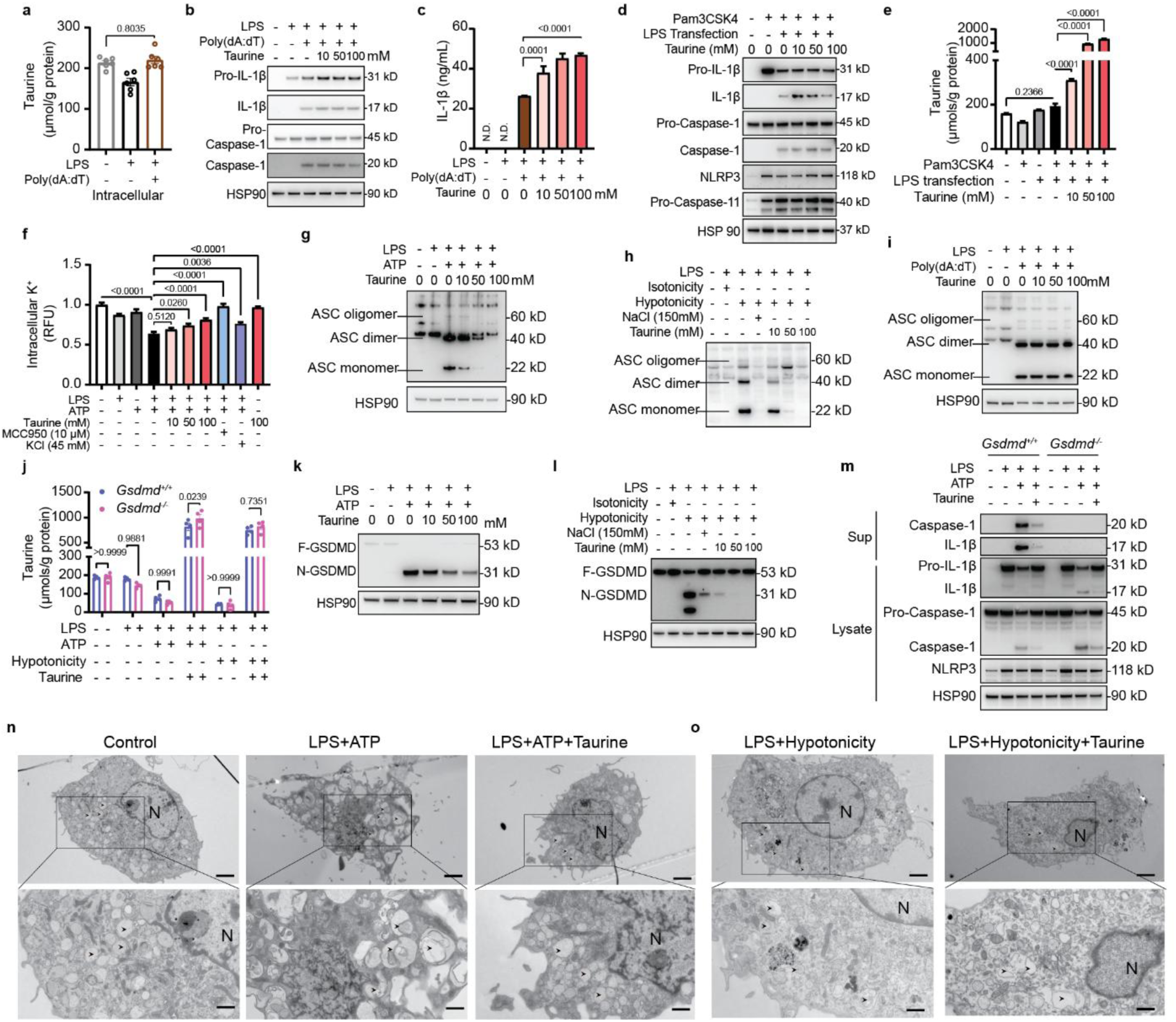
Taurine inhibits NLRP3 inflammasome assembly and GSDMD activation. **(a-c)** LPS-primed BMDMs are transfected with Poly(dA:dT) in the presence of taurine. (a) Intracellular taurine levels of BMDMs. (b) Western blots analyses of IL-1β (p17), caspase-1 (p20) in cell lysates and supernatants of AIM2 inflammasome activated BMDMs. (c) Production of IL-1β from these BMDMs, measured by ELISA. These results are representative of three to five independent experiments. **(d)** Immunoblots of cell lysates and supernatant from BMDMs stimulated with Pam3CSK4 and transfected with LPS in the presence of taurine. Results are representative of three independent experiments. **(e)** Intracellular taurine levels of BMDMs primed with Pam3CSK4 and transfected with LPS in the presence of taurine. **(f)** Quantification of intracellular potassium following inflammasome activation with LPS and ATP in the presence of indicated concentration of taurine. Intracellular potassium was detected by potassium indicator, ION Potassium Green-2 AM. **(g)** Immunoblot of ASC in cell lysates and cross-linked cytosolic insoluble pellets from BMDMs stimulated with LPS and ATP with taurine. Results are representative of three independent experiments. **(h)** Western blots detection of ASC oligomerization in cross-linked insoluble pellets from BMDMs activated by hypotonic solution in the presence of taurine or sodium chloride. Results are representative of three experiments. **(i)** Immunoblot of ASC oligomerization in BMDM stimulated with LPS and transfected with poly(dA:dT) in the presence of taurine. These results are representative of three independent experiments. **(j)** Intracellular taurine levels in control and *Gsdmd^-/-^* BMDMs primed with LPS and treated with ATP, hypotonicity. **(k)** Immunoblot of cell lysates from BMDM activated by LPS and ATP in presence of taurine. F-GSDMD, full length GSDMD; N-GSDMD, N-terminal GSDMD; C-GSDMD, C-terminal GSDMD. The results are representative of five experiments. **(l)** Immunblot detection of GSDMD cleavage in BMDM activated by LPS and hypotonicity in presence of taurine. **(m)** Western blots of IL-1β (p17), caspase-1 (p20) in cell lysates and supernatants of control and *Gsdmd^-/-^* BMDMs stimulated with LPS and ATP with taurine. These results are representative of three independent experiments. **(n-o)** Transmission electron micrograph of LPS-primed BMDMs and treated with ATP (n) or hypotonicity (o). **(n)** Resting macrophages containing autophagosomes with clear membrane structure (left). In response to LPS and ATP stimulation, macrophages containing autophagosomes with apparent remnants of degenerating parts (middle). With taurine addition, autophagosome with clear autophagosome vacuoles (right). **(o)** Hypotonic solution induced cell swelling in LPS-primed BMDMs (left) and in presence of taurine. Scale bar: 2 μm in lower magnification (1,900 ×); 1 μm in higher magnification (4800 ×). Arrows point to the autophagosome. N, nucleus.

It is known that upstream K^+^ efflux is a trigger common to many NLRP3 activators ^42^ for inflammasome assembly. Interestingly, we found that taurine dose-dependently prevent K^+^ efflux triggered by the LPS and ATP (Fig. 3f). To determine the requirement of K^+^ efflux step in taurine mediated inflammasome regulation, we activated the macrophages with imiquimod, a K^+^ independent activator of NLRP3 inflammasome ^47^. Notably, taurine inhibited imiquimod induced IL-1β while restoration of K^+^ with KCl had minor effect on reducing IL-1β (Fig. S4a, S4b). Imiquimod also caused approx. 38% taurine efflux and restoration of taurine inhibited imiquimod induced caspase-1 cleavage (Fig. S4b, S4c). Inhibition of NLRP3 by MCC950 and KCl did not rescue taurine efflux from imiquimod activated macrophages (Fig. S4c). Next, we utilized NLRP3 activator Nigericin which is a specific K^+^/H^+^ antiporter. Nigericin which acts specifically at K^+^ channel to cause K^+^ efflux causes approx. 20% taurine efflux (Fig. S4d). This efflux of taurine in LPS+Nigericin activated cells was not rescued by MCC950, validating prior results of taurine in upstream regulation of NLRP3 inflammasome activation (Fig. S4d). In presence of taurine, nigericin induced caspase-1 cleavage and IL-1β release was reduced (Fig S4e, f). In addition to TLR4, taurine also inhibits TLR1/2 stimulator Pam3CSK4 licensed inflammasome activation in response to ATP (Fig. S4g, S4h) and nigericin (Fig. S4i, S4j). These data further suggest a model whereby taurine responds to danger as an upstream event to induce downstream NLRP3 inflammasome activation (Fig. S4k).

Next, we examined NLRP3 inflammasome assembly through formation of NLRP3-dependent ASC oligomers, a key event that nucleates ASC for inflammasome activation^3–5^. Cytosolic fractions from cell lysates were cross-linked, and ASC monomers and higher order complexes were observed after stimulation with LPS and ATP. Taurine inhibited the ASC-complex formation in response to NLRP3 activation in macrophages (Fig. 3g). Notably, in TLR4 primed macrophages, hypotonicity induced ASC oligomerization was also inhibited by taurine supplementation (Fig. 3h). Confirming specificity, taurine had no effect on AIM2 inflammasome activation (LPS and Poly(dA:dT) induced ASC-complex formation) (Fig. 3i).

Given taurine inhibited NLRP3 inflammasome, we next investigated pyroptosis, a caspase-1 dependent lytic cell death ^48^ that is driven by cleavage of gasdermin D (GSDMD) and activation of its pore-forming domain ^49^. It is thought that nonselective ion flux through GSDMD pores causes the collapse of the macrophage electrochemical gradient leading to lytic cell death ^49^. However, extracellular ATP and hypotonicity induced taurine efflux is independent of GSDMD pore (Fig. 3j). Consistent with upstream role of taurine in inhibiting NLRP3 induced caspase-1 activation, taurine led to dose dependent inhibition of GSDMD cleavage in macrophages activated by extracellular ATP (Fig. 3k), hypotonic solution (Fig. 3l), urate crystals (Fig. S5a) and ceramide C6 (Fig. S5b). Consistent with no effect on caspase-11 inflammasome, taurine treatment did not impact Pam3CSK-primed and intracellular LPS transfection induced GSDMD cleavage (Fig. S5c). In support of prior results, taurine inhibited GSDMD activation in LPS primed, imiquimod (Fig. S5d) and nigericin (Fig. S5e) activated macrophages. And taurine suppressed TLR1/2 primed, ATP (Fig. S5f) and nigericin (Fig. S5g) activated macrophages. Furthermore, consistent with upstream role of caspase-1 in cleavage of full length GSDMD to N-truncated active GSDMD, taurine’s inhibitory effects on caspase-1 by NLRP3-induced activation was maintained in *Gsdmd* deficient macrophages which had equivalent increase in caspase-1 cleavage by extracellular ATP (Fig. S5h, Fig. 3m) and hypotonicity (Fig. S5i, S5j).

Given taurine’s strong inhibitory effects on inflammasome and gasdermin-D mediated pyroptosis, we next conducted transmission electron microscopy to determine ultrastructural changes in macrophages activated in response to NLRP3 activators and hypotonicity. Compared to control cells, LPS and ATP treated cells displayed classic features of a pyroptotic activation response, with loss of pseudopodia and accumulation of large fused autophagosomes. Interestingly, taurine protected against the formation of large autophagosomes and pyroptosis (Fig. 3n, Fig. S6). Similarly, hypotonicity-induced cell swelling was reduced in presence of taurine (Fig. 3o, Fig. S7). Further TEM analyses and quantification revealed that taurine maintained mitochondrial structural integrity in inflammasome activated macrophages (Fig. S8a, S8b, S8c). Taken together, these data demonstrate that taurine inhibits NLRP3 activation by acting upstream and blocking K^+^ efflux, ASC oligomerization, and GSDMD mediated pyroptosis.

## Taurine inhibits NLRP3 in models of cryopyrinopathy independently of Slc6a6 and Slc36a1

Missense mutations in NLRP3 cause human systemic inflammatory diseases like familial cold autoinflammatory syndrome (FCAS) and Muckle–Wells syndrome (MWS), which are characterized by overproduction of IL-1β and IL-18 ^50^. We tested the efficiency of taurine in BMDMs with *Nlrp3* mutations (L351P and A350V) corresponding to the human NLRP3 gain-of-function mutations (L353P and A352V), resulting in human FCAS and MWS respectively, rendering the inflammasome constitutively active without the requirement of NLRP3 ligands ^50^. In mouse macrophages with FCAS mutation that renders NLRP3 constitutively active, taurine blocked the processing of both IL-1β and caspase-1 (Fig. 4a, 4b). The GSDMD cleavage was inhibited in the presence of taurine in response to LPS in FCAS BMDMs (Fig. 4c). Similarly, taurine can also suppress IL-1β release (Fig. 4d), caspase-1 processing (Fig. 4e) and GSDMD activation (Fig. 4f) in TLR4 primed mouse macrophages with the *Nlrp3* mutation that mimics MWS (Fig. 4d-f). These data suggest that elevation of taurine in macrophages inhibits intrinsic NLRP3 inflammasome assembly in disease models of cryopyrinopathies.

**Fig. 4.**
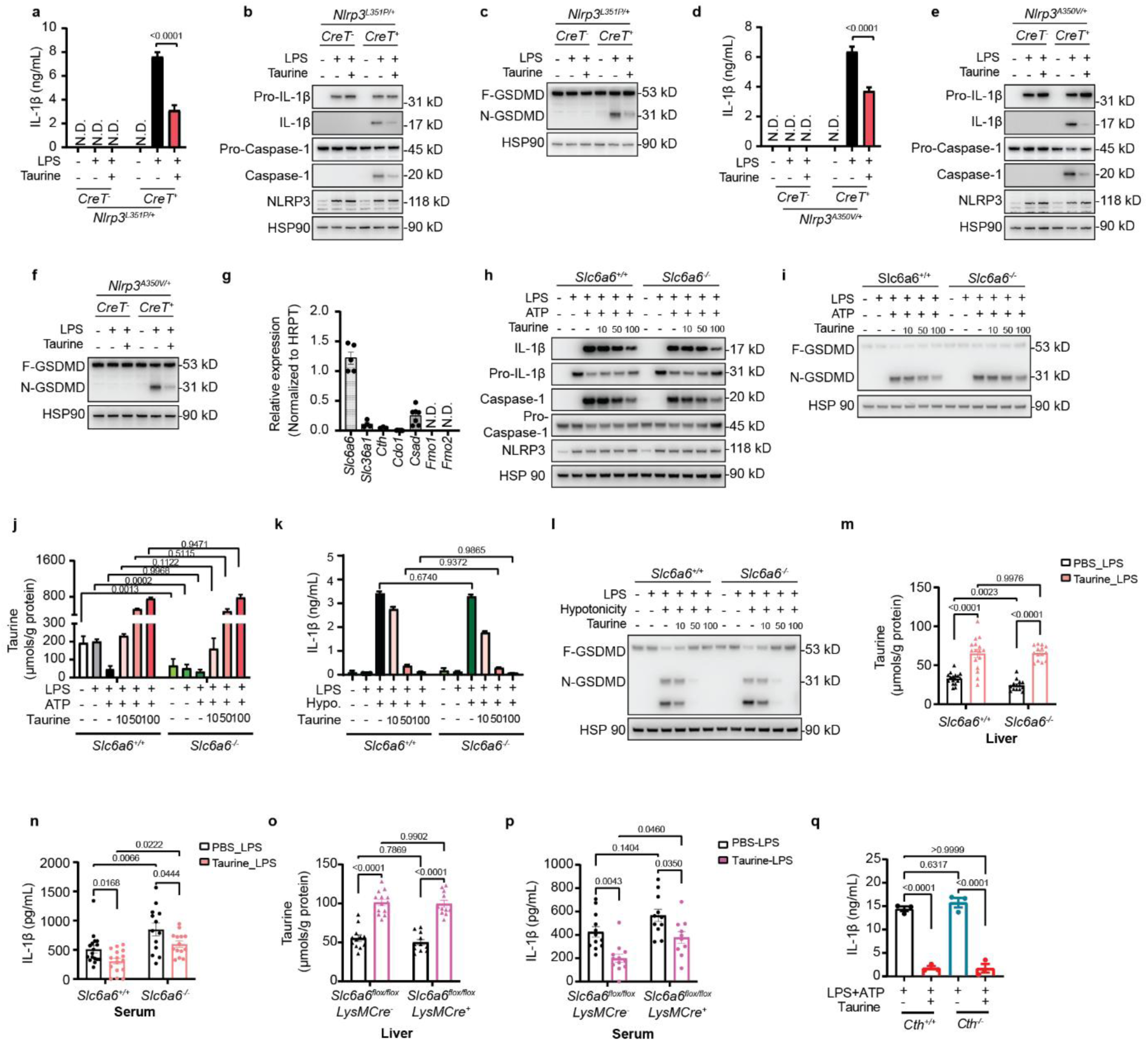
Taurine inhibits NLRP3 in disease models of cryopyrinopathies independently of Slc6a6 or Slc36a1 transporters. **(a-c)** BMDMs from ***Nlrp3^L^***^351^***^P^*** mice. **(a)** IL-1β production from these BMDMs stimulated with LPS and treated with taurine as measured by ELISA. These data are representative of three independent experiments carried out in triplicate. **(b)** Western blots of IL-1β (p17), caspase-1 (p20) in cell lysates and supernatants from BMDMs stimulated with LPS and treated with taurine. These results are representative of three independent experiments. **(c)** Immunoblot analysis of GSDMD cleavage in cell lysates from BMDM stimulated with LPS and treated with taurine. These results are representative of three independent experiments. **(d-f)** BMDMs from *Nlrp3^A^*^350^*^V^* mice. (d) IL-1β production from BMDMs stimulated with LPS and treated with taurine. These data are representative of three independent experiments carried out in triplicate. (e) Western blots of IL-1β (p17), caspase-1 (p20) in BMDMs stimulated with LPS and treated with taurine. These results are representative of three independent experiments. (f) Western blot analyses of GSDMD cleavage in BMDM stimulated with LPS and treated with taurine. These results are representative of three independent experiments. **(g)** qPCR analysis of genes encoding key proteins involved in taurine uptake and synthesis in BMDMs (n=5-6). **(h-i)** Western blot analyses of IL-1β (p17), caspase-1 (p20) (h) and GSDMD (i) from control and *Slc6a6^-/-^* BMDMs stimulated with LPS and ATP and treated with taurine. These results are representative of three independent experiments. **(j)** Intracellular taurine levels in control and *Slc6a6^-/-^* BMDMs activated by LPS and ATP and treated with taurine. **(k-l)** IL-1β production (k) and GSDMD cleavage (l) in control and *Slc6a6^-/-^* BMDMs stimulated with LPS and hypotonicity and treated with taurine. These data are representative of three independent experiments carried out in triplicate. **(m and n)** 10-to 11-week-old littermate *Slc6a6^+/+^* and *Slc6a6^-/-^* mice were pretreated with taurine (1000 mg/kg body weight) or vehicle control for 5 days, then given LPS i.p. injection, 4 hours later blood and liver were analyzed (n=17,16,14,14). (m) Taurine levels of the liver from the mice. (n) Serum levels of IL-1β from these mice. **(o and p)** 13- to 14-week-old littermate *Slc6a6*^flox/flox^*LysMCre*^-^ or *Slc6a6*^flox/flox^*LysMCre*^+^ mice were pre-treated with taurine (1000 mg/kg body weight) or vehicle control for 5 days, IL-1β was quantified in blood and liver 4 h post LPS challenge (n=12,12,10,10). Taurine levels in livers (o) and serum IL-1β levels (p) of control and myeloid specific Slc6a6 deficient mice. **(q)** Production of IL-1β from *Cth^+/+^*and *Cth^-/-^* BMDMs stimulated with LPS and ATP and treated with taurine. Data are expressed as the mean ± SEM of three independent experiments carried out in triplicate.

Taurine is zwitterionic and its high concentration in specific cells could be due to either synthesis downstream of cysteine or by active import via canonical transporters Slc6a6 and Slc36a1 ^51^. We found that murine BMDMs had low expression of *Cdo1*, *Csad* with undetectable expression of *Fmo1* and *Fmo2* (Fig. 4g). In contrast, BMDM had high expression of taurine transporter *Slc6a6* compared to *Slc36a1* (Fig. 4g). As Slc6a6 is reported to be the main taurine transporter in cells ^52,53^, we reasoned that Slc6a6 might transport taurine in macrophages. Genetic ablation of *Slc6a6* led to about 80% depletion of the taurine in serum (Fig. S9a). However, the *Slc6a6^-/-^*BMDMs treated with taurine showed comparable IL-1β (Fig. S9b), caspase-1 (Fig. 4h) and GSDMD cleavage (Fig. 4i) in response to LPS and ATP treatment. Furthermore, taurine caused a dose-dependent reduction of LDH release (Fig. S9c) in *Slc6a6^-/-^* macrophages. Compared to control cells, the *Slc6a6* deficient macrophages displayed equivalent reduction of ASC oligomerization (Fig. S9d) in response to taurine. Furthermore, the *Slc6a6* deficient macrophages showed efficient taurine efflux and import upon NLRP3 activation (Fig. 4j). Taurine also inhibited the hypotonicity induced NLRP3 inflammasome activation (Fig. 4k, Fig. S9e) and GSDMD cleavage (Fig. 4l) independently of Slc6a6, since the deficiency of Slc6a6 has no effect on intracellular taurine uptake (Fig. S9f). We next silenced *Slc36a1* (*Pat1*) in BMDM to investigate the potential role of this transporter in taurine’s effects on macrophage biology (Fig. S9g). Compared to Slc6a6, alternate taurine transporter Slc36a1 has low mRNA and protein expression in BMDMs (Fig. 4g, Fig. S9h, S9i). Taurine significantly and equivalently inhibited IL-1β release in both control and Slc36a1 silenced BMDMs (Fig. S9j). To rule out potential compensation of one transporter, we created *Slc6a6* and *Slc36a1* double deficient BMDMs. Interestingly, compared to control BMDMs, taurine equivalently inhibited LPS and ATP induced IL-1β secretion in BMDMs lacking both canonical taurine transporters *Slc6a6* and *Slc36a1* (Fig. S9k). The livers of *Slc6a6* deficient mice treated with LPS displayed lower baseline taurine (Fig. 4m) but with equivalent reduction of IL-1β by taurine treatment (Fig. 4n). Given inflammasome activation requires TLR priming, there was no spontaneous activation of NLRP3 in *Slc6a6* deficient mice with lower baseline taurine. We next conditionally ablated Slc6a6 in myeloid cells and found no significant change in taurine levels in liver (Fig. 4o) and both control littermates and LysMCre:Slca6^fl/fl^ mice had equivalent reduction of serum IL-1β upon taurine supplement in response to LPS challenge (Fig. 4p) validating our results from *in vitro* cell model.

The enzyme cystathionine-γ lyase (CTH) mediates the critical rate limiting step in TSP downstream of cysteine to generate taurine (Fig. 1b) ^16^. Given our finding that CR induced elevation of taurine is associated with reduction of cysteine despite an increase in *Cth* expression (Fig. 1a, 1b), we investigated the role of CTH in taurine’s effect on macrophages. When compared control to *Cth* deficient macrophages, taurine similarly inhibited the NLRP3 inflammasome dependent caspase-1 cleavage (Fig. S9l) and IL-1β secretion (Fig. 4q). Consistent with our data that inflammasome activation does not change oxidized cystine or GSH in macrophages (Fig. 2b, Fig. S2d), taurine inhibited IL-1β (Fig. S9m), caspase-1 and GSDMD activation in macrophages independently of redox regulator NRF2 (Fig. S9n). This suggest that inflammasome deactivation is independent of taurine synthesis and taurine uptake from media is sufficient to inhibit caspase-1 cleavage, IL-1β release and GSDMD mediated pyroptosis in macrophages without the requirement of canonical transporters Slc6a6 and Slc36a1.

## Elevation of taurine inhibits inflammaging in mice and enhances healthspan

We next investigated the effects of taurine on inflammation *in vivo*. Adult mice were pre-treated for 7 days with taurine and were assessed 4 hours after LPS challenge (Fig. 5a). Pretreatment with taurine significantly reduced serum concentrations of IL-1β, TNFα and MCP-1 (Fig. 5a) without affecting IL-6 (Fig. S10a). Unlike LPS, urate crystals specifically activate NLRP3. We found that taurine treatment resulted in the decreased production of GRO-α/KC and MCP-1 (Fig. S10b) but not TNF-α or IL-6 (Fig. S10c) in response to *in vivo* challenge with MSU crystals. Taurine also inhibited LPS induced IL-1β concentration in female mice (Fig. 5b). Acute administration of taurine (4 hours) in LPS+ATP treated mice also reduced IL-1β in adult mice (Fig. 5c) without affecting the body weight (Fig. 5d). We next administered lethal dose of LPS to mice to induce inflammation and sepsis. Interestingly consistent with its anti-inflammatory mechanism of action, the taurine treatment significantly reduced endotoxemia induced mortality in adult mice (Fig. 5e). Given taurine levels increase with age and inhibit the NLRP3 inflammasome, we next investigated the mechanism of taurine in control of age-related inflammation. The 20-month-old C57BL/6 mice were treated with taurine in drinking water for 4 months and evaluated for measures of healthspan and inflammation. Taurine treatment doubled taurine concentrations in the serum (Fig. 5f) and liver of old mice (Fig. S10d). Taurine supplementation reduced the body weight and fat mass of old mice without affecting the lean mass (Fig. S10e, S10f). Consistent with lower body weight, the old mice treated with taurine had significant improvements in the glucose homeostasis (Fig. 5g). Notably, taurine also improved rotarod test and grip strength, which suggested enhanced balance, motor coordination and reduced frailty, which are important indicators of healthspan (Fig. 5h, 5i).

**Fig. 5.**
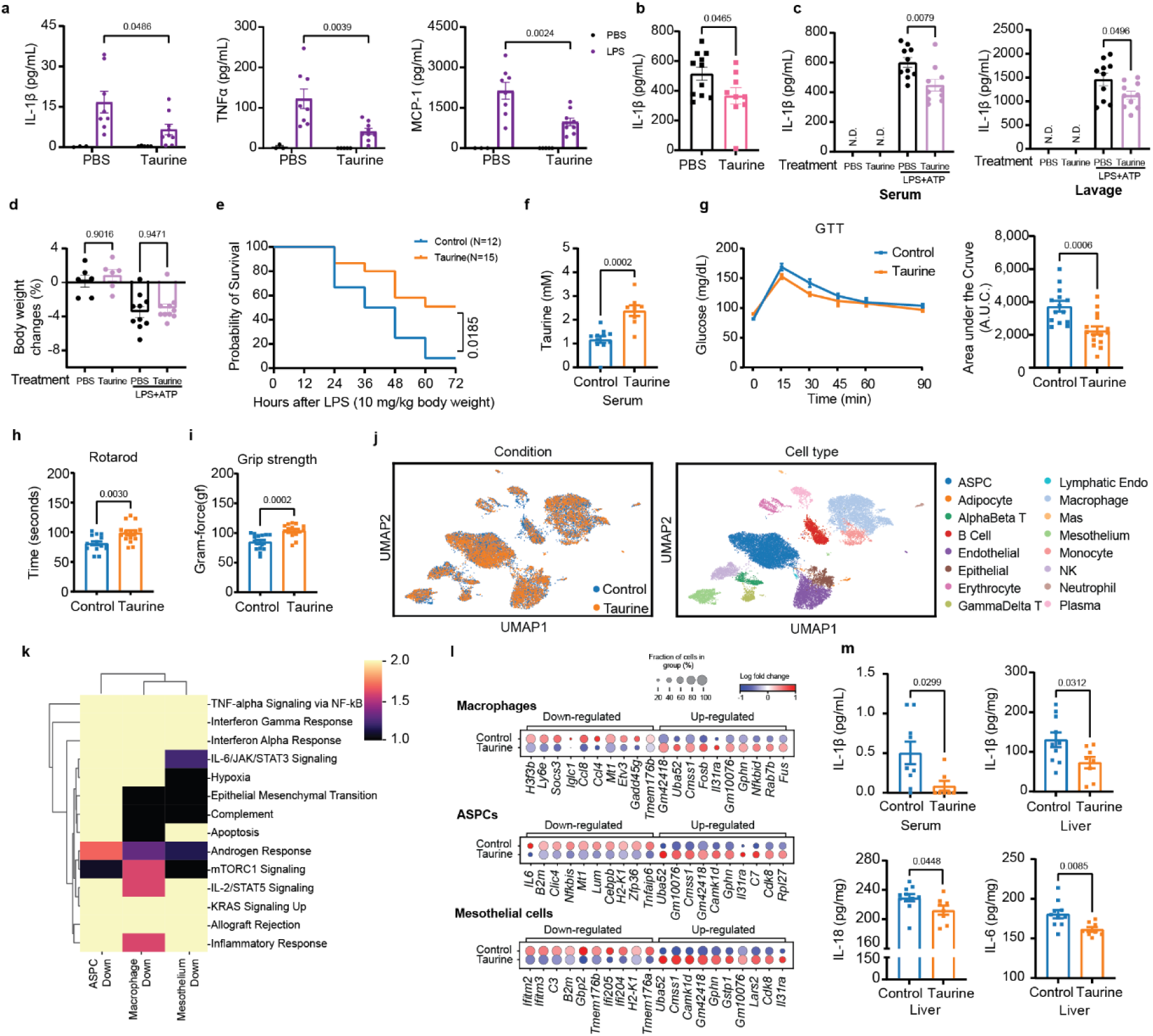
Elevation of taurine inhibits NLRP3-inflammaging and enhances healthspan. **(a)** 8- to 9-week-old C57BL/6J male mice were intraperitoneally injected with taurine at 500 mg/kg body weight or vehicle control for 7 days and then give i.p. LPS injection. 4 hours later, serum levels of IL-1β, TNF-α, and MCP-1 were measured by ELISA (n=3, 8, 4, 9) **(b)**10- to 11-week-old C57BL/6J female mice were i.p. injected with taurine at 1000 mg/kg body weight or vehicle control for 5 days, and then mice were given i.p. LPS injection for 4 hours. Serum IL-1β was measured by ELISA (n=11, 9). **(c and d)** 13-week-old C57BL/6J male mice pretreated with taurine (1000 mg/kg body weight) or vehicle control for 5 days, then mice were given LPS for 4 hours followed by ATP treatment for 15 mins (n=6,6,10,10). (c) Levels of IL-1β of the serum and peritoneal lavage were measured by ELISA. (d) body weight changes of these mice. **(e)** Survival curves of 10- to 11-week-old C57BL/6J male mice pretreated with PBS or Taurine (1000 mg/kg body weight) for 5 days followed by LPS i.p. injection (n=12,15). **(f-m)** Aged male C57BL/6N mice (20-month-old) given normal water or drinking water with taurine (8,000 mg/kg/day, 4% in water (w/v)) for 4 months. **(f)** Serum taurine (n=11,8) levels after 4 months of oral taurine treatment. **(g)** Glucose tolerance test (GTT) and area of the curve (A.O.C.) of mice for 3 months taurine treatment (n=14, 15). **(h and i)** Rotarod (h) and Grip-strength (i) tests of mice after 3 month of taurine treatment (n=15, 15). **(j)** UMAP visualizations of the scRNA-sequencing data from stromal vascular faction (SVF) cells of visceral adipose tissue (VAT) in these old mice (n=10, 8 pooled). Identified cells colored by treatment condition (left). Proportions and annotation of selected cell types in control versus taurine treated condition (right). **(k)** The mouse hallmark gene set enrichment analysis for down-regulated genes in taurine treated group across cell types. Color represents minus log10 adjusted p-values. **(l)** Dot plots showing significantly up/down-regulated genes (n=10 each) in taurine-treated group versus control group for macrophages, ASPCs and mesothelial cells. The dot colors indicate the log fold changes. Dot sizes indicate the fraction of cells that express the genes. **(m)** Serum levels of IL-1β and liver levels of IL-1β, IL-18 and IL-6 from mice treated with taurine for 4 months (n=9, 8). Error bars represent the mean ± SEM.

To determine the mechanism of taurine’s anti-inflammaging effects, we conducted single-cell RNA (sc-RNA) sequencing of stromal vascular fraction cells isolated from visceral adipose tissue (VAT) of old mice that were supplemented with taurine. The uniform manifold approximation and projection (UMAP) unbiased clustering revealed 16 distinct cell populations including adaptive, innate immune cells and mesothelial cells (Fig. 5j). Comparison of sham with taurine groups revealed few changes in cellular composition including no change in adipose progenitor cells (ASPCs), macrophages or T cell except reduction in B cells, which are known to increase in VAT of aged WT mice ^12^ (Fig. 5j, Fig. S10g). The gene set enrichment analysis (GSEA) analyses within ASPC, macrophages and mesothelial cells displayed significant downregulation of pro-inflammatory pathways, including tumor necrosis factor (TNF)-α signaling, interferon-response and IL-6 signaling and complement cascade (Fig. 5k). The taurine induced top up and down regulated genes in ASPCs, macrophages and mesothelial cells displayed common gene-regulation hubs such as upregulation of *Uba52* and *Cmss1* and inhibition of β2 microglobulin (*B2m*) expression (Fig. 5l, Fig. S10h). Previous studies have shown that increase of B2M, a component of major histocompatibility complex (MHC) class I, is a pro-aging factor that is linked to inflammation and cognitive decline in old mice ^54^. Collectively, these data suggested that taurine downregulates major pro-inflammatory pathways in aged mice. To confirm this, we performed cytokines detection in serum and liver tissues in these aged mice. Compared to sham-treated mice, supplementation of taurine significantly decreased the concentrations of IL-1β (both serum and liver), IL-18 and IL-6 (liver) (Fig. 5m), the pro-inflammatory cytokines that drive inflammaging. Taurine also has an inhibitory trend in IL-18 in the serum without affecting Monocyte chemoattractant protein-1 (MCP-1) and eotaxin (Fig. S10i). Collectively these data demonstrate that taurine downregulates NLRP3 inflammasome and major pro-inflammatory pathways in aged mice.

## Discussion

Activation of TSP pathway, that generates cysteine and culminates in production of GSH and taurine has previously been associated with pro-longevity interventions ^22,55^. In addition, taurine has been linked to protection from inflammation in SARS-COV2 infection and reduces oxidative stress ^56,57^. In older adults, chronic inflammation contributes to loss of organ function and development of aging-associated diseases ^1–3^. Our findings indicate that CR-induced upregulation of taurine in humans is an adaptive mechanism to control inflammation. Reduction of taurine was reported to be a driver of aging, and daily gavage of old mice for approximately two years with taurine enhanced lifespan^21^. Our data demonstrate that taurine concentrations do not decline and instead increase in older humans and mice. The reasons for the difference in findings are unclear, one possibility is that taurine is also present in high concentrations in red blood cells, leukocytes, so that hemolyzed plasma or serum or contamination with degranulated platelets or other cells could influence taurine measurements. Our sample sets were carefully evaluated for this potential confounder, and we can rule out that increased taurine is a result of cell contamination. It has been known that taurine levels increase in response to osmolar imbalance, thirst or increased hypertonicity ^16^. Notably, aging is associated with reduced thirst ^58^, which can induce negative water balance and lead to adaptive increase in organic osmolytes such as taurine. Indeed, using isotope-tracking ^2^H methods that quantifies water turnover through measurement of hydrogen isotope dilution and elimination, aging in humans was found to be associated with reduced water turnover ^59^. Thus, the observed age-related increase in taurine could also represent an adaptive compensatory response to maintain osmolar balance.

Our findings that inflammasome activation causes taurine efflux in macrophages and pyroptosis is consistent with prior reports of swelling-evoked taurine release in the brain, including astrocytes and neurons ^60,61^. In addition, metabolomics identified greater taurine efflux in macrophages undergoing pyroptosis than apoptosis ^62^. Our results provide direct evidence that taurine efflux acts as an upstream sensor of DAMPs that activates downstream NLRP3 inflammasome in TLR primed macrophages. Taurine did not affect the priming step, its efflux induces K^+^ release, ASC oligomerization, caspase-1 activation and GSDMD cleavage. Taurine was active against the hypersensitive *Nlrp3* mutations associated with FCAS and MWS, and against canonical NLRP3 stimuli without impacting AIM2, NLRC4 or caspase-11 inflammasomes. Taurine functions in mM ranges in multiple organs and upon efflux by NLRP3 activating DAMPS while 10-50 mM supplementation effectively inhibits the inflammasome. Thus, unlike endocrine hormones or a distally produced metabolite that have to be secreted in blood to produce an effector response, taurine’s mM cellular and local concentrations likely operate in autocrine/paracrine fashion without undergoing the dilution effect in blood. Consistent with Mangum and Towle’s hypothesis, elevation of taurine may serve as an enantiostatic metabolite that opposes inflammaging to maintain biological function despite perturbation of internal milieu.

Given its low lipophilicity, taurine is impermeable across plasma membrane and high levels of cellular taurine are maintained by synthesis and transport via Slc6a6. Our studies performed in *Cth^-/-^* macrophages have shown that synthesized taurine has no effect on NLRP3 inflammasome activation. Surprisingly, canonical taurine transporters were not required for anti-inflammasome effect in macrophages as both the global and myeloid cell specific *Slc6a6* deficient mice had a similar anti-inflammatory response to taurine upon LPS challenge and NLRP3 activation. Interestingly, macrophages can take up intact mitochondria and multiple cellular components including exosomes via unknown mechanisms ^63^. It has been shown that taurine can boost cellular uptake of small D-peptides by endocytosis or trogocytosis ^64^. In some condition, uptake of taurine appeared to be non-saturable and possibly due to diffusion ^65^. Our data suggest that additional mechanisms are required for taurine uptake in macrophages and future studies are required to determine the transport and efflux mechanism.

One to two years of mild food restriction leading to reduced calorie intake in healthy adults in CALERIE-II clinical trial helped highlight the transsulfuration pathways (TSP), which is typically dormant in fed conditions with adequate protein intake. This is likely because reduction of food consumption is associated with proportional decrease in protein intake and humans prioritize production of sulfonic amino acid taurine by rewiring cysteine metabolism in TSP. Upon exposure to NLRP3 activating ‘danger signals’ taurine efflux allows downstream NLRP3 inflammasome assembly while elevation of taurine inhibits NLRP3 driven inflammation and inflammaging. In summary, we demonstrate that elevation of taurine in aging and CR-induced metabolic stress maybe an adaptive mechanism to restore homeostasis. Our studies suggest that elevation of taurine inhibits age-related inflammation by serving as a metabolic osmosensor that protects against NLRP3 inflammasome activation.

## Acknowledgments

We thank all investigators and staff involved in coordinating and executing CALERIE-II clinical trial. We also thank to Genentech for providing anti-Caspase-1 antibody. SR is a recipient of American Federation for Aging Research (AFAR) postdoctoral fellowship. The research in Dixit Lab was supported in part by NIH grants AG031797, AG045712, P01AG051459, AR070811, and Cure Alzheimer’s Fund (CAF).

## Author Contributions

CG conducted majority of experiments, designed the studies, did data analysis and participated in writing the manuscript. SR conducted the taurine supplementation studies in old mice and performed the healthspan analyses. LO and HHK performed RNA-sequencing data analyses of CALERIE-II samples. YHY and TG performed mouse phenotyping. SM and ACS generated pre and post vaccination human samples and assisted in data analyses. MD and YK performed sc-RNA-sequencing analysis, data analyses, statistical tests and data interpretation. ER and SRS were involved in the design of parent CALERIE-II trial, participant recruitment and execution of study at PBRC-Baton Rouge clinical site. SRS also performed the biopsies of human adipose tissue. JO and HMH contributed the cryopyrinopathy models and in experimental design to generate macrophages mimicking human MWS and FCAS diseases. RM and YS performed the metabolomic analyses of human adipose tissue and conducted the data analyses. All authors participated in manuscript preparation. VDD conceived the project, helped with data interpretation, analysis and wrote the manuscript.

## Materials and Methods

Supplementary Text Figs. S1 to S10

## Materials and Methods

### Human

#### Human Calorie Restriction Study

This study used samples from participants of CALERIE Phase 2 study. CALERIE Phase 2 is a multi-center, parallel group, randomized controlled trial recruiting non-obese healthy individuals. 238 metabolically healthy adults (BMI: 22.0-27.9 kg/m^2^, Age: 25-45) were enrolled in the study from Pennington Biomedical Research Center (PBRC, Baton Rouge, LA), Washington University (St. Louis, MO) and Tufts University (Boston, MA) (NCT00427193), Duke University, (DCRI; Duke coordinating research initiative Durham, NC) worked as a coordinating center. Study participants were randomized assigned to the 25% CR intervention versus ad-libitum control group who continued their habitual diet. Men were between 20 and 50 years old and women were between 20 and 47 years old. Their body mass index (BMI) was between 22.0 and 27.9 kg/m^2^ at the initial visit. The diet intervention was designed to achieve 25% CR, defined as a 25% reduction from ad-libitum baseline energy intake. CR group participants accomplished 14% of CR. 14 participants who completed 1 year of intervention provided abdominal subcutaneous adipose tissue by biopsy and plasma that were subjected to metabolomics analysis. Only 8 participants completed all three timepoints baseline, 1- and 2-years CR. All participants that provided adipose tissue at all three timepoints after CR were studied for gene expression changes. All studies were performed under the protocol approved by the Pennington institutional review board with the informed consent from participants.

#### Human Blood Samples

Serum samples were obtained from a previous study of young (age 21-35) and older (age ≥ 65) adults^66^ who were recruited during the 2018-2019 influenza season according to an approved protocol by the Human Subjects Research Protection Program of the Yale School of Medicine. Informed consent was obtained from all participants prior to sample collection. Participants reporting acute illness two weeks prior to recruitment were not included in the study. The exclusion criteria included subjects with history of malignancy, cirrhosis, subjects undergoing hemodialysis, subjects using immunomodulating medications as well as subjects with primary or acquired immune deficiency. All subjects received the high seasonal influenza vaccine (Fluzone High-Dose) containing hemagglutinin (HA) proteins from A/Michigan/45/2015 X-275 (H1N1), A/Singapore/INFIMH-16-0019/2016 IVR-186 (H3N2), and B/Maryland/15/2016 BX-69A (a B/Colorado/6/2017-like virus, B Victoria lineage) at a dose of 60 μg for each HA. Blood was collected via venipuncture in a BD Vacutainer serum tube at the indicated timepoints prior to and post-vaccination. The tubes were allowed to clot at room temperature for 2 hours before centrifugation at 2500 RPM for 20 min. The clear serum as supernatant was stored in 500 μl aliquots at -80°C.

#### Mice

Mice in Fig. 5 and Fig. S10 (long term taurine supplement) were from NIA on C57BL/6N background, all other wild-type mice were on C57BL/6J genetic background. C57BL/6J mice were purchased from Jackson Laboratory and maintained in our lab. *Nlrp3^-/-^* mice (B6.129S6-*Nlrp3*^tm1Bhk^/J, Strain# 021302) were purchased from Jackson Laboratory and maintained in our lab. *Cth^-/-^* mice (C57BL/6NTac-Cth^tm1a^(EUCOMM)^Hmgu/Ieg^) were purchased from the European Mouse Mutant Cell Repository. *Slc6a6^-/-^*mice (C57BL/6NCrl-*Slc6a6*^em1^(IMPC)^Mbp^) were purchased from the Mutant Mouse Resource & Research Centers supported by NIH). *Slc6a6^flox/flox^*mice (C57BL/6JGpt-Slc6a6^em1Cflox^/Gpt, strain ID: T019182) were purchased from GemPharmatech, China. LysM Cre mice (B6.129P2-Lyz2^tm1^(cre)^Ifo^/J, Strain #: 004781) were purchased from The Jackson Laboratory. *Nlrp3*^L351P^ gain of function familial cold autoinflammatory syndrome (FCAS) and *Nlrp3*^A350V^ Muckle–Wells syndrome (MWS) knock-in mutations have been described before ^50,51^. Briefly, the *Nlrp3 (L351P^neoR/+)^* and *Nlrp3 (A350V^PneoR/+)^* mutation were bread to a tamoxifen inducible Cre (Cre/ESR1) and conditionally activated in vitro by treating cells with 4-hydroxy tamoxifen (Sigma, 1 μM, for 24 h). The *Gsdmd^-/-^* mice were provided by Thirumala-Devi Kanneganti from St. Jude Children’s Research Hospital. The *Nrf2^-/-^* mice were provided by Andrew Wang from Yale University. Animals were maintained under specific pathogen-free facility in the Yale Animal Resource Center. For breeding cages, we used Teklad Global 19% Protein Extruded Rodent Diet (Sterilizable)_2019S (no taurine is this diet). Weanlings were put on standard laboratory chow (PicoLab Laboratory Rodent Diet_5L0D*) ad libitum. This diet contains 0.03% taurine. All animal procedures were ethically approved by the Institutional Animal Care and Use Committee (IACUC) at Yale University prior to experimentation.

#### Generation of Murine BMDMs

8-12-week-old mice were euthanized by isoflurane inhalation and death was confirmed by cervical dislocation. Bone marrow was subsequently harvested from femurs and cells were differentiated in RPMI (Gibco) containing L929 cell supernatant (30%), fetal bovine serum (20%, from Thomas Scientific), M-CSF (10ng/ml, R&D System) and Antibacterial/Antimycotic (1%, Thermo Fisher Scientific) for 6 days, after which cells were counted and plated at 2×10^6^ cells/ well in 6-well plate.

#### Mice Treated with Taurine (orally)

Mice received standard chow (PicoLab Laboratory Rodent Diet_5L0D*) ad libitum. Taurine was orally administered via water at dose level of 0 (Control) or 8,000 mg/kg/day for 4 months in 20-month-old wild type male mice. We speculated that administering taurine orally via water should be effective since 4 months of taurine supplemented drinking water induces about 2-fold increase of serum taurine concentration. Amount of water intake per mouse was calculated by measuring the water consumption every day for each cage.

#### Body Composition Analyze

Mice were subject to magnetic resonance imaging machine (EchoMRI; Echo Medical Systems) to measure body composition and quantify fat mass and lean mass. Fat mass % and lean mass % were calculated based on total mass.

#### Glucose Tolerance Tests

For glucose tolerance testing, mice were fasted overnight (∼16 hours) with baseline fasting blood glucose level recorded as 0-min glucose. Then glucose (0.4 g/kg body weight, Sigma) was intraperitoneally injected in awake mice using an insulin syringe. Blood glucose at 15, 30 45, 60, and 90 min timepoints by Contour Next EZ Blood Glucose Monitoring system.

#### Rotarod Test

A rotarod (Med-Associates, St Albans, VT) apparatus was used to assess motor function and coordination. The machine consists of a slowly accelerating plastic rod and five testing lanes. The rod is 3.2 cm in diameter, supported 30 cm above the base of the apparatus. Before the test day, mice were trained on the Rotarod for 3 days. Holding the mouse by the tail, each mouse was place on the rotating rod, facing away from the direction of rotation so it must walk forward to stay upright. The rotarod was accelerated slowly up to a specified rpm and not exceeding 40 rpm. Mice were monitored and the length of time the animal stayed on the rotarod were recorded. Mice were given 3 trials per day, with an inter-trial interval of 30-40 min. Each trial lasted a maximum of 300 s.

#### Grip Strength Measurement

The Grip Strength test is used to evaluate motor function as maximal muscle strength of forelimbs. The mouse’s forepaws were placed on a wire grid, while its tail was gently pulled backwards. The maximum strength of the grip prior to grip release was recorded. The test was performed in three sessions per week with each session consisting of three trials. The average performance for each session was presented as the average of the three trials.

#### In Vivo LPS Challenge

8- to 9-week-old male C57BL/6J mice were intraperitoneally injected daily with Taurine (500 mg/kg body weight, dissolved in sterile PBS or PBS alone (same volume of taurine solution) for 7 days before LPS challenge. At 7 days, mice were injected intraperitoneally with LPS (2mg/kg body weight, Sigma) suspended in sterile PBS for 4 hours. 4 hours later mice were euthanized, and the serum concentrations of IL-1β, MCP-1, TNF-α and IL-6 were measured by Luminex (Thermo Fisher Scientific).

10- to 11-week-old female C57BL/6J mice were intraperitoneally injected daily with taurine (1000 mg/kg body eight, dissolved in sterile PBS or PBS alone (same volume of taurine solution) for 5 days before LPS challenge. At day 5, mice were injected intraperitoneally with LPS (2mg/kg body weight) suspended in sterile PBS for 4 hours. 4 hours late mice were euthanized, and the blood was collected. The serum concentration of IL-1β was measured by ELISA (R&D).

#### MSU-Induced Peritonitis Model

2-month-old male C57BL/6J male mice were intraperitoneally injected with Taurine (500mg/kg body weight, dissolved in sterile PBS) or PBS (same volume of taurine solution) only daily for 7 days. At 7 days, mice were injected intraperitoneally with PBS or MSU (2.5 mg/mouse, Invivogen) suspended in PBS for 4 hours. 4 hours later mice were euthanized, and the serum concentrations of GRO-α, MCP-1, TNF-α and IL-6 were measured by Luminex (Thermo Fisher Scientific).

#### Acute in vivo LPS + ATP challenge

Thioglycolate was intraperitoneally injected for inducing in vivo model of peritoneal macrophages. 4% thioglycolate 1 mL was injected once 4 days before LPS and ATP treatment. 4 days later, 13-week-old C57BL/6J male mice were intraperitoneally injected with 4% Taurine (1000mg/kg body weight) and LPS (2 mg/kg body weight) at same time, 4 hours later mice were intraperitoneally injected with ATP (1 mM/kg body weight). 15 mins later mice were euthanized, blood and peritoneal lavage were collected. The body weight was recorded before treatment and at the time of sacrifice. The concentrations of IL-1β were measured by ELISA (R&D).

#### Peritoneal lavage fluid collection

Fill a 5 mL syringe with cold Opti-MEM medium. With a 20-G needle, insert needle through peritoneal wall and inject 5 mL of the Opti-MEM medium into each mouse. Using the same syringe and needle, aspire fluid from peritoneum. Centrifuge 10 min at 400 g, 4°C. The supernatant (lavage fluid) was used to do the following ELISA IL-1β test. The peritoneal macrophages were separated from the cell pellet by using the macrophage isolation kit (Peritoneum).

#### Peritoneal macrophage isolation

Macrophage isolation (peritoneum) was performed following the manufacturer’s instruction (Miltenyi Biotec 130-110-434). Collect cell suspension as before. Pass cells through 40 μm filter to remove cell clumps. Then magnetic labeling with the antibody cocktail. Then harvest the peritoneal macrophages by magnetic separation with MS Columns.

#### LPS-lethal Survival Rates

Following treatment of mice with PBS or Taurine for 5 days. Then the mice were i.p. with LPS (10 mg/kg body weight). The survival rates were determined daily, every 12 h, for 4 days.

#### Canonical NLRP3 inflammasome Assays

For stimulation, 2×10^6^ differentiated BMDMs were plated overnight, the medium was changed to opti-MEM the following morning, then the cells were primed with 1 μg/ml LPS (Sigma)or Pam3CSK4 (1 μg/ml, Invivogen) with or without taurine (Sigma) for 4 hours. Primed cells were then treated with 5 mM ATP (Sigma) for 45 min, 250 μg/ml MSU (Invitrogen) for 6 hours or 100μM Ceramide C6 (Cayman) for 6 hours, Nigericin (10 μM, Invivogen, 1h) or Imiquimod (50 μM, Invitrogen, 2h) to activate the NLRP3 inflammasome. MCC950 (10 μM, Sigma) or KCl was added 15 min before ATP or Nigericin treatment. The treatments with glycine (Sigma), glutamine (Sigma), and folic acid (Thermo Fisher Scientific) were administered at same time with LPS.

#### Reducing Media Osmolarity Induced NLRP3 Inflammasome Activation

LPS-primed BMDMs were stimulated with isotonic solution (PBS, pH 7.4) and hypotonic solution (dilute isotonic solution 1:4 with ultrapure sterile water) for 1 hour to induce NLRP3 inflammasome activation.

#### AIM2 Inflammasome Assays

In experiments to activate the AIM2 inflammasome, LPS-primed cells were transfected with 2 ng/μl poly(dA:dT) (Invitrogen) using Lipofectamine 2000 Transfection Reagent (Thermo Fisher Scientific) for 6 hour following the manufacturer’s instruction.

#### NLRC4 inflammasome assays

In experiments to activate the NLRC4 inflammasome, LPS-primed cells were transfected with 500 µg/ml flagellin (Invivogen tlrl-epstfla)) using DOTAP Liposomal Transfection Reagent (Sigma 11202375001) for 6 hour following the manufacturer’s instruction.

#### Taurine Assessment

Treated BMDMs were washed with PBS for at least 3 times to rule out the culture medium taurine contamination. 2×10^6^ BMDMs were resuspended in 100 μl PBS with protease inhibitor cocktail (Sigma) using a cell scraper, then cells were homogenized by passing through an insulin needle 20 times and spun at 14,000 rpm for 10 min at 4 °C. 25 μl lysate was transferred to a 96-well plate and then incubated with taurine detecting reagent mix (Abcam or Cell Biolabs) for 30 min at 30 °C. Measure absorbance in an endpoint mode by using a microplate reader.

Intracellular concentration is normalized to macrophages protein level.

For serum taurine assessment, 10-25 μl serum was used to incubate with taurine detecting reagent mix and followed by the manufacturer’s instructions (Abcam).

#### Non-canonical Inflammasome assays

Bone marrow cells were differentiated for 5-7days. Adherent BMDMs cultured overnight in 6-well plates at 1 × 10^6^ cells per ml, total 2ml, before being primed for 5–6 h with 1 μg ml^−1^ Pam3CSK4 in OPTI-MEM (Life Technologies). Primed cells were transfected with 2 μg ml^−1^ LPS by using 0.25% v/v FuGENE HD (Promega) for 10 h to active the non-canonical inflammasome.

#### Western Blotting

Supernatant was removed from BMDMs (2×10^6^) following stimulation and lysates were harvested in 100μl RIPA buffer (Sigma) with protease inhibitor (Sigma). The lysates were then centrifuged at 14,000 rpm for 10 min, supernatant was collected, and the protein concentration was measured using the DC Protein Assay (Bio-Rad). Samples were mixed with NuPAGE Sample Reducing Agent (Invitrogen) and NuPAGE LDS Sample buffer (Life technologies). Mixed samples were boiled at 95 °C for 10 min prior to loading into a SDS-PAGE gel (Invitrogen). Same amounts of protein (20-30 μg) and supernatant (40 μl) were loaded on a SDS-PAGE gel. The Invitrogen gel running system was used to resolve protein and Bio-Rad semi-dry transfer system was used for electrophoretic transfer of proteins onto nitrocellulose membrane. Following transfer, the membrane was incubated in milk (5% in TBST, Bio-Rad) for 1 hour and subsequently incubated in primary antibodies rolling overnight at 4 °C. Host-matched secondary antibodies were incubated with membrane for 1 hour at room temperature. The ECL substrate (Thermo Fisher Scientific) was used to detect bands by using a ChemiDoc MPTM Imaging System (Bio-Rad). The primary antibodies used were anti-IL-1β (GeneTex), anti-Caspase-1 (p20) (mouse), mAb (Casper-1) (Adipogen), anti-GSDMD (Abcam), anti-NLRP3/NALP3, mAb(Cryo-2) (Adipogen), anti-Asc, pAb(AL177) (Adipogen), anti-HSP 90 (Cell Signaling Technology), Phospho-IκBα (Ser32/36) (5A5) Mouse mAb (Cell Signaling Technology), IκBα Antibody (Cell Signaling Technology), Phospho-NF-κB p65 (Ser536) (93H1) Rabbit mAb (Cell Signaling Technology) , NF-κB p65 (D14E12) XP® Rabbit mAb (Cell Signaling Technology), Phospho-p44/42 MAPK (Erk1/2) (Thr202/Tyr204) Antibody (Cell Signaling Technology), p44/42 MAPK (Erk1/2) (137F5) Rabbit mAb (Cell Signaling Technology), Phospho-SAPK/JNK (Thr183/Tyr185) (81E11) Rabbit mAb (Cell Signaling Technology), SAPK/JNK Antibody (Cell Signaling Technology), Phospho-p38 MAPK (Thr180/Tyr182) (D3F9) XP® Rabbit mAb (Cell Signaling Technology), and p38 MAPK (D13E1) XP® Rabbit mAb (Cell Signaling Technology), Slc36a1 polyclonal antibody (Proteintech), Slc36a1 polyclonal antibody (Invitrogen). The secondary antibodies used were Goat anti-Rabbit IgG(H+L) Secondary Antibody, HRP (Invitrogen) and Goat anti-mouse IgG (H&L) Secondary Antibody, HRP (Thermo Fisher Scientific).

#### ASC Oligomerization

BMDMs were stimulated as before, the cells were rinsed with ice-cold PBS, and 500 μl ice-cold NP-40 lysis buffer (20 mM HEPES-KOH, pH 7.5, 150 mM KCl, 1 % NP40, 0.1 mM PMSF and protease inhibitor cocktail) was added. Cells were left on ice for 15 min followed by centrifugation at 6,000 rpm at 4 °C for 10 min, then the supernatant was collected for immunoblot analysis. The lysate was washed once in NP-40 lysis buffer and resuspended in 50 μl NP-40 lysis buffer with 2mM Disuccinimidyl suberate (Sigma), which were incubated at room temperature for 30 min with rotation. Samples were then centrifuged, and the cross-linked pellets were re-suspended in 20 μl SDS sample buffer. Samples were boiled for 5 min at 95 °C and analyzed by Western blotting.

#### ELISA and Cytokine Profiling by Multiplex Cytokine Assay

Blood was collected and incubated at room temperature for 30 min, centrifuge at 3,000 rpm at 4 °C for 20 min to isolate serum. The concentrations of IL-1β, IL-18, IL-6 and TNFα of supernatant from stimulated BMDMs and serum were determined by ELISA (R&D system) or 28 plex magnetic bead panel Luminex (Invitrogen) following the manufacturer’s instructions with appropriate dilution. Absorbance at 450 nm was then quantified using a Thermo Fisher plate reader. Corrected absorbance values were calculated by subtracting the back-ground absorbance (540nm).

#### LDH Assay

The LDH Assay Kit (Cytotoxicity) (Abcam) was used to quantify lactate dehydrogenase release from cells as a measure of cell death in BMDMs following inflammasome stimulation. Freshly harvested supernatants were detected in this assay following the manufacturer’s instructions. The percentage was compared with the positive control (100%).

#### Transmission Electron Microscopy Analysis

Differentiated BMDMs were seeded on chamber slides (Thermo) overnight, and stimulated on second day. The the cells on coverslips were Fixed in 2.5% glutaraldehyde in 0.1 M Sodium cacodylate buffer, pH 7.4 followed by rinsing three times in 0.1 M sodium cacodylate buffer. The cells were then post-fixed in 0.5% Osmium tetroxide reduced with 0.8% Potassium ferrocyanide in 0.1 M Sodium cacodylate buffer, pH 7.4. The cells were rinsed three times in 0.1 M sodium cacodylate buffer followed by rinsing with HPLC water. For en bloc staining, the cells were incubated in 2% Uranyl acetate for 45 min followed by rinsing in HPLC water. The cells were dehydrated in an 50%, 70%, 90%, and 100% ethanol series and embedded with EPON epoxy resin. The sample blocks were sectioned with a UC7 Ultracut Ultramicrotome (Leica) and 60 nm thick sections were laid on formvar-coated Nickel mesh grids (Electron Microscopy Sciences). The grids were post-stained with 2% Uranyl acetate and Reynold’s Lead citrate. Images were acquired on an 80 kV BioTwin Transmission electron microscope using an AMT NanoSprint15 MKII camera.

#### Analyzing Transmission Electron Microscopy with ImageJ

The mitochondrial morphology, length and width were analyzed by ImageJ. Track the outer mitochondrial membrane of each mitochondrion. Draw the straight line along the major and minor axis of each mitochondrion, click the measure function to obtain mitochondrial area, lengths and widths. ∼300 mitochondrion from 26 cells (to average about 10 mitochondria per cell) were measured.

Analyzing mitochondrial cristae, split each image into four quadrants and pick the left two quadrants to analyze, use the same quadrants for all images. Trace the outline of each crista within the mitochondrion. Click the measure function to obtain the area of each cristae, and cristae surface is the sum of the area of all the cristae in a single mitochondrion. To determine the cristae number, count the number of cristae in each mitochondrion.

#### Intracellular Potassium Assay

Intracellular potassium was detected by K+ indicator, ION Potassium Green-2 AM (Abcam, ab142806). 1×10^5^ BMDMs/ well were seeded into a 96 well plate and then treated with NLRP3 inflammasome stimulators. For the NLRP3 inflammasome activated BMDMs, ION potassium green-2 AM (ab142806; Abcam) was added to a final concentration of 5 μM and cells were incubated for a further 2 hours in the dark at room temperature. Cells were washed twice with cold PBS to remove excess fluorescent dye. Then 50ul PBS was added to the stained cells in the wells. Ion K+ green fluorescence was captured on a fluorescence plate reader (Ex/Em= 510/546 nm).

#### Cell transfection

The siRNA sequences of Slc36a1 are from Thermo Fisher Scientific (Cat# 4390771). Oligonucleotides were transfected into the indicated cells using the Lipofectamine RNAiMAX transfection reagents, siRNAs against *Slc36a1* were transfected into target cells sequentially at an interval of 48 h.

#### RT-qPCR

Total RNA was extracted from the indicated cells and then reverse-transcribed into cDNA using a iScript cDNA synthesis kit (Bio-Rad). qPCR was performed on the Thermo ABI QuantStudio 7 Flex Real-Time PCR with a SYBR Green Universal Master Mix (Thermo Fisher Scientific). The data were normalized to the expression of HRPT.

#### Sample preparation for metabolome analysis

Frozen tissues or serum samples, together with internal standard compounds (mentioned below), were subjected to sonication in 500 μL of ice-cold methanol. To this, an equal volume of ultrapure water (LC/MS grade, Wako, Japan) and 0.4 volume of chloroform were added. The resulting suspension was centrifuged at 15,000 × g for 15 min at 4 °C. The aqueous phase was then filtered using an ultrafiltration tube (Ultrafree MC-PLHCC, Human Metabolome Technologies, Japan), and the filtrate was concentrated by nitrogen spraying (aluminum block bath with nitrogen gas spraying system, DTU-1BN/EN1-36, TAITEC, Japan). The concentrated filtrate was dissolved in 50 μL of ultrapure water and utilized for IC-MS and LC-MS/MS analysis. Methionine sulfone and 2-morpholinoethanesulfonic acid were employed as internal standards for cationic and anionic metabolites, respectively. The recovery rate (%) of the standards in each sample measurement was calculated to correct for the loss of endogenous metabolites during sample preparation.

#### IC-MS metabolome analysis

Anionic metabolites were detected using an orbitrap-type MS (Q-Exactive focus; Thermo Fisher Scientific, USA) connected to a high-performance ion-chromatography (IC) system (ICS-5000+, Thermo Fisher Scientific, USA) that allows for highly selective and sensitive metabolite quantification through IC separation and Fourier transfer MS principle. The IC system included a modified Thermo Scientific Dionex AERS 500 anion electrolytic suppressor, which converted the potassium hydroxide gradient into pure water before the sample entered the mass spectrometer. Separation was carried out using a Thermo Scientific Dionex IonPac AS11-HC column with a particle size of 4 μm. The IC flow rate was 0.25 mL/min, supplemented post-column with a makeup flow of 0.18 mL/min MeOH. The potassium hydroxide gradient conditions for IC separation were as follows: from 1 mM to 100 mM (0–40 min), to 100 mM (40–50 min), and to 1 mM (50.1–60 min), with a column temperature of 30 °C. The Q Exactive focus mass spectrometer was operated in the ESI-negative mode for all detections. A full mass scan (m/z 70–900) was performed at a resolution of 70,000. The automatic gain control target was set at 3 × 10^6 ions, and the maximum ion injection time was 100 ms. The source ionization parameters were optimized with a spray voltage of 3 kV, and other parameters were as follows: transfer temperature, 320 °C; S-Lens level = 50, heater temperature, 300 °C; sheath gas = 36, and Aux gas, 10.

#### LC-MS/MS metabolome analysis

Cationic metabolites were measured using liquid chromatography-tandem mass spectrometry (LC-MS/MS). The LCMS-8060 triple-quadrupole mass spectrometer (Shimadzu corporation, Japan) with an electrospray ionization (ESI) ion source was employed to perform multiple reaction monitoring (MRM) in positive and negative ESI modes. The samples were separated on a Discovery HS F5-3 column (2.1 mm I.D. x 150 mm L, 3μm particle, Sigma-Aldrich) using a step gradient of mobile phase A (0.1% formate) and mobile phase B (0.1% acetonitrile) with varying ratios: 100:0 (0-5 min), 75:25 (5-11 min), 65:35 (11-15 min), 5:95 (15-20 min), and 100:0 (20-25 min). The flow rate was set at 0.25 mL/min, and the column temperature was maintained at 40°C.

#### Bulk RNA-Sequencing and Analysis

Subcutaneous adipose tissue collected using biopsy of CALERIE phase 2 study participants was used for whole tissue RNA sequencing. The RNA was extracted using Zymo miniprep kits. RNA that passed quality checks was used for RNA sequencing library preparation at Yale Center by using HiSeq 2500.

The sequencing quality of raw reads was assessed with FastQC (v0.11.3). Raw reads were mapped to hg38 human genome (GENCODE v29) (26) using STAR aligner (v2.5.3a) (27) with the following options: --outFilterMultimapNmax 15 --outFilterMismatchNmax 6 --outReadsUnmapped Fastx -- outSAMstrandField intronMotif --outSAMtype BAMSortedByCoordinate. Quality control of the mapped reads was done using Picard tools (v2.18.4). Quantification was done using htseq-count function from HTSeq framework (v0.9.1): htseqcount–f bam –r pos –s no –t exon.

#### scRNA sequence

20-month-aged male mice were given taurine drinking water (Taurine group) or normal water (Control group) for 4 months, 8 mice from control group and 10 mice from taurine group were sacrificed to collect visceral adipose tissue (VAT). VAT was digested in Hanks Buffered Salt Solution (Thermo Fisher Scientific) including 0.1% collagenase II enzyme (Worthington Biochemicals) for 40 min in shaking water bath at 37 °C. Red blood cells were lysed with ACK lysing buffer (Thermo Fisher Scientific) and cells were passed through 40 μm filters, pooled samples per group (control n=8, taurine n=10), 10,000 cells were analyzed using Chromium Next GEM Automated Single Cell 3’ cDNA Kit v3.1(10X genomics).

The count matrices were concatenated and converted into AnnData format [1]. We filtered out possible doublet with number of genes per cell larger than 8000. We further filtered out low-quality cells with mitochondria reads larger than 15% of the total counts. Then we performed library size normalization and log1p transformation by Scanpy ^67^ with the default settings. Then we selected top 1000 highly variable genes by Scanpy function pp, highly_variable_genes, with all other settings set to default. The highly variable genes normalized expression were used to compute the 50 leading principal components by Scanpy. We run leiden clustering and UMAP visualization by Scanpy using default settings. We next annotated Leiden clusters’ cell type assignment using marker genes from ^68^. The dataset was annotated into 16 cell types, including adipocytes, adipose stem and progenitor cells (ASPCs), vascular cells and immune cells. Two small Leiden clusters with cell numbers 57 and 18 were dropped due to the expression of epididymal markers (Spink8) and high droplet likelihood by co-expression of macrophage and NK cell markers. The final dataset comprises 21571 non-zero expressed genes and 19904 cells. 12107 of the 19904 cells are from the control condition, and 7797 of the 19904 cells are from the taurine treated condition.

The differential expression analysis for each cell type was performed by Wilcoxon rank-sum test implemented in Scanpy (tl.rank_gene_groups). Genes with adjusted p-values smaller than 0.05 were selected to be the differential expressed genes for each cell type. Then we performed the pathway enrichment analysis for each cell type by GSEAPy ^69^ with the mouse hallmark gene set_70_.

