## Supplemental Figures for "Hormetic elevation of taurine restrains inflammaging by deactivating the NLRP3 inflammasome"

Fig. S1_Revision 1_02242025

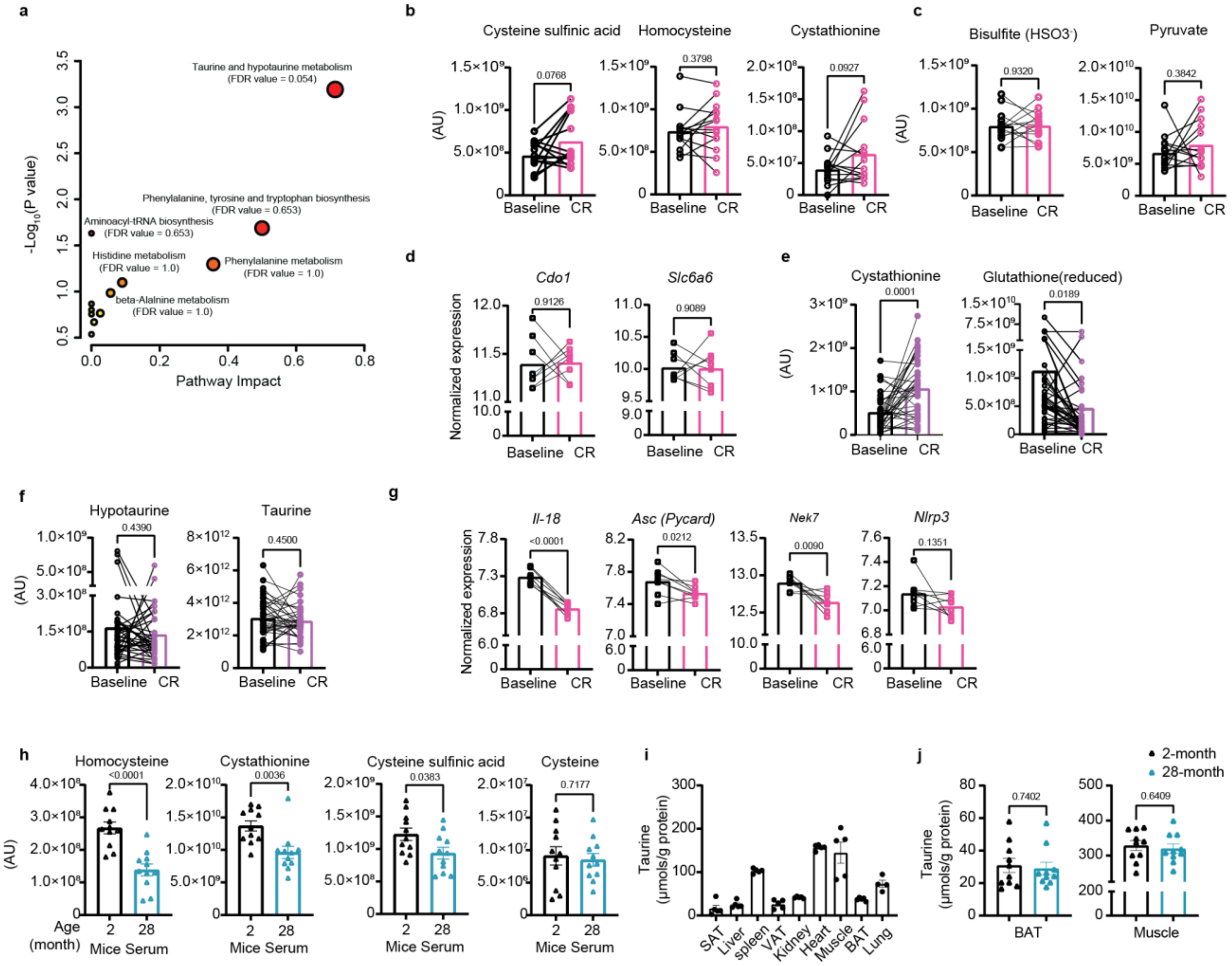


**Fig. S1. Caloric restriction in human adipose tissue shifts transsulfuration pathway from GSH to taurine metabolism.**

**(a)** Enrichment analysis was conducted using MetaboAnalyst (ver 5.0) with subcutaneous adipose tissue (SAT) metabolites significantly upregulated following CR compared to baseline. **(b-c)** Quantification of cystathionine, homocysteine, cysteine sulfinate (b) and bisulfite ($HSO_3^-$), pyruvate (c) by LC-MS/MS in human SAT after 1 year CR (n=14). **(d)** RNA-Sequencing analysis of the expression levels of *Cdo1* and *Slc6a6* after 1 year CR in SAT (n=8). **(e)** Changes in cystathionine and reduced glutathione measured by LC-MS/MS in human plasma after 1 year CR (n=37). **(f)** Quantification of hypotaurine and taurine by LC-MS/MS in human plasma after 1 year CR. Statistical differences were calculated by using paired t-tests (n=37). **(g)** The determination of gene expression of *Il-18*, *Asc*, *Nek7* and *Nlrp3* by RNA sequencing in human SAT at baseline and one year after CR. (**h**) Serum concentration of homocysteine, cystathionine, cysteine sulfinate and cysteine measured by LC-MS/MS in 2-month-old and 28-month-old male C57/B6N mice (n=11/group). **(i)** Determination of taurine concentration in various tissues in C57/B6J mice (n=4-5). **(j)** Taurine concentrations of mice muscle and BAT from 2-month-old and 28-month-old C57/B6J male mice (n=10/group). Error bars represent the mean ± SEM.

Fig. S2_Revision 1_02242025

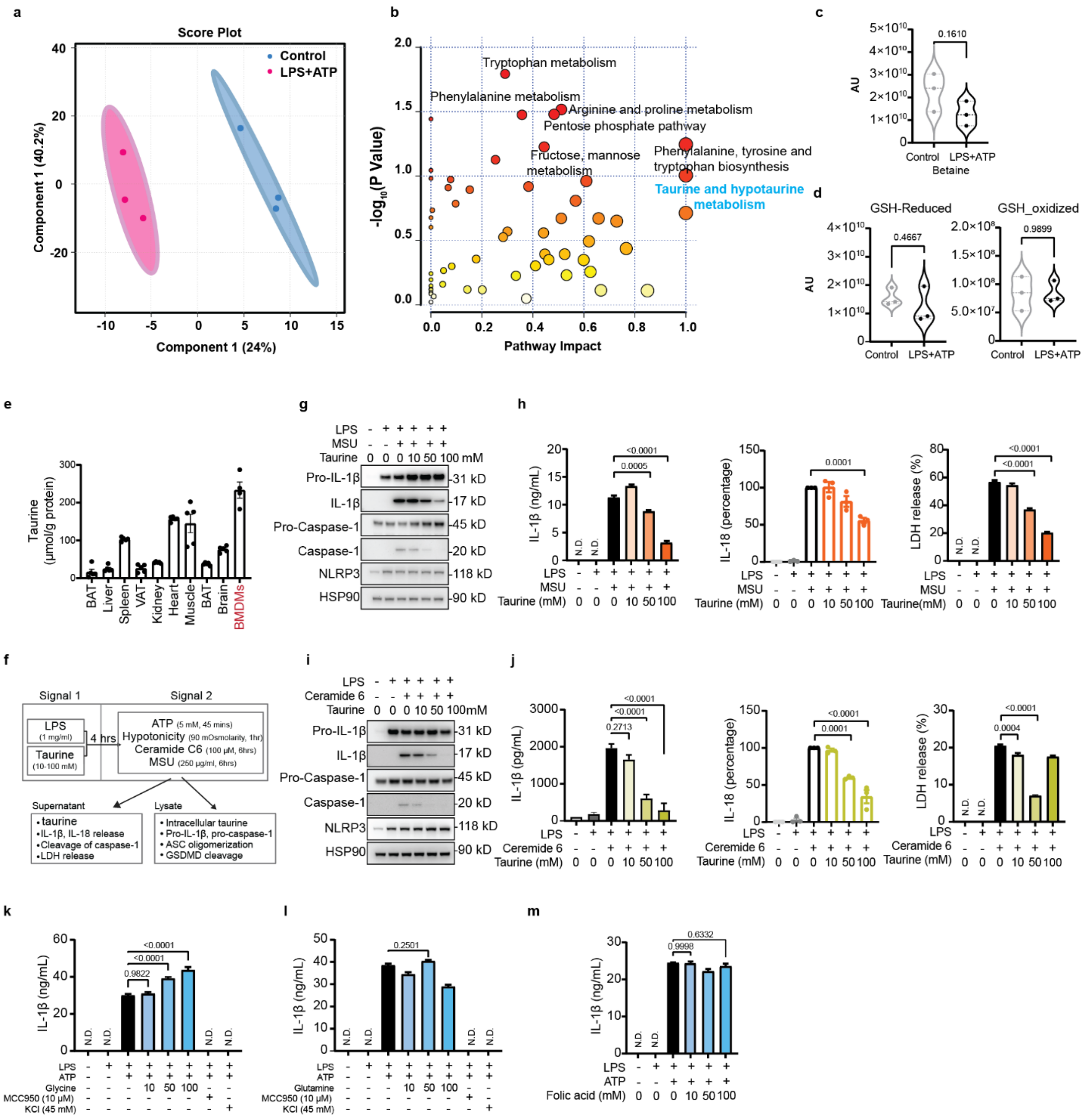


**Fig.S2. Taurine inhibits NLRP3 inflammasome activated by ‘danger signals’ urate and ceramide.**

**(a)** Pathway analysis (targeted) was conducted using MetaboAnalyst with metabolites significantly regulated following LPS and ATP treatment compared to Control. **(b)** Scores plot of partial least squares-discriminant analysis (PLS-DA) based on metabolomics data between LPS and ATP treated BMDMs and control BMMDs (n=3). Shaded circles represent 95% confidence intervals, while colored dots illustrate individual samples. The axes are labelled by the first and second components with the percentages of variance of the data explained by that component in parentheses. **(c-d)** Changes in the intracellular betaine (c), reduced GSH and oxidized GSH (d) in response to LPS and ATP stimulation in BMDMs, measured by MS/MS. **(e)** Taurine concentration in various tissues and BMDMs isolated from adult C57/B6J mice (n=4-5)**. (f)** Experimental design of NLRP3 inflammasome assay to evaluate the impact of taurine on BMDMs. **(g)** Western blots of IL-1β and caspase-1 in cell lysates and concentrated supernatants from BMDMs stimulated with LPS and MSU and treated with taurine. These results are representative of three independent experiments. **(h)** Production of IL-1β, IL-18 and LDH release from BMDMs primed with LPS, and stimulated with MSU in presence of taurine. **(i)** Western blots of IL-1β and caspase-1 in cell lysates and concentrated supernatants from BMDMs stimulated with LPS and Ceramide 6 and treated with taurine. These results are representative of three independent experiments. **(j)** Production of IL-1β, IL-18 and LDH from BMDMs primed with LPS and stimulated with Ceramide 6 in presence of taurine. (These results are representative of three independent experiments. **(k-m)** ELISA analysis of IL-1β production from BMDMs stimulated with LPS (1 μg/ml, 4 hours), followed by ATP (5 mM, 45 min) treatment with or without glycine (k), glutamine (l) or folic acid (m) supplementation. Error bars represent the mean ± SEM.

Fig. S3_Revision 1_02242025

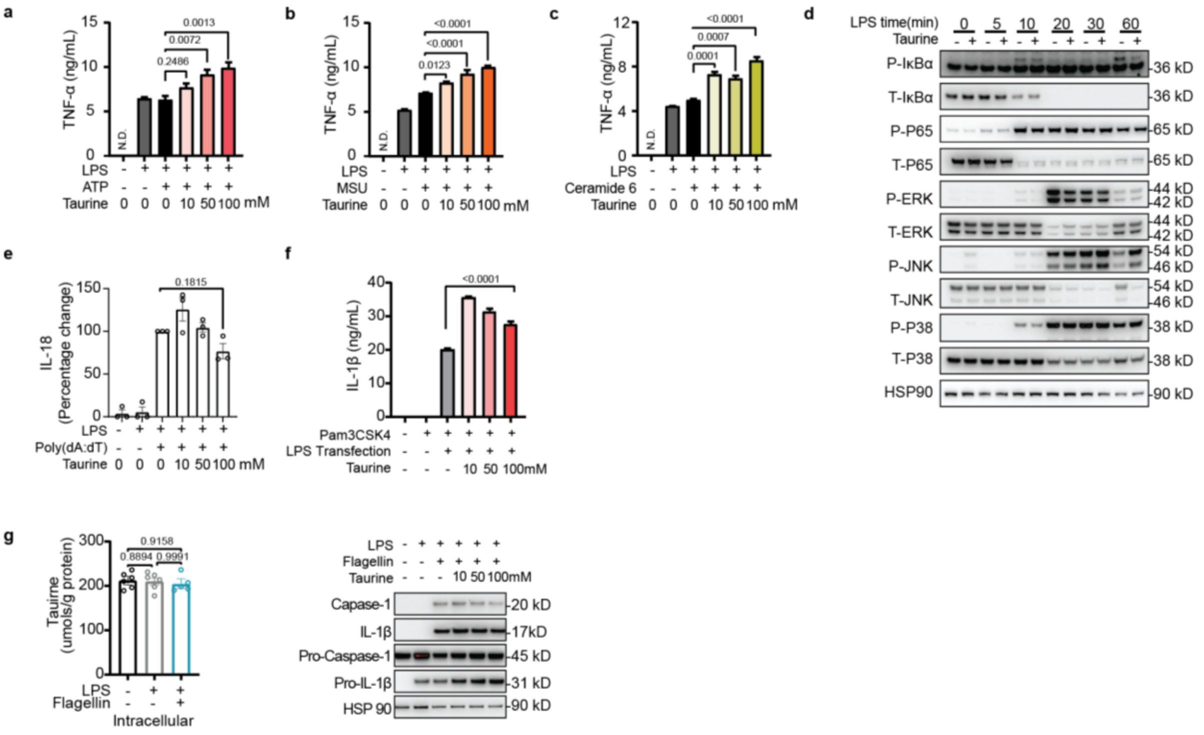


**Fig.S3. Taurine does not impact NLRP3 inflammasome priming.**

**(a-c)** Production of TNFα (as measured by ELISA) from BMDMs stimulated with LPS and ATP (a), MSU (b), Ceramide C6 (c) and treated with taurine. These data are representative of five independent experiments carried out in triplicate. **(d)** Western blots of cell lysates from BMDM stimulated by LPS and taurine. (P: Phosphorylation; T: Total). **(e)** Production of IL-18 (as measured by ELISA) from BMDMs stimulated with LPS and transfected poly(dA:dT) and treated with taurine, percentage compared with LPS and Poly(dA:dT) treatment. **(f)** Production of IL-1β from BMDMs stimulated with Pam3CSK4 and transfected with LPS and treated with taurine as measured by ELISA. These data are representative of five independent experiments carried out in triplicate. **(g)** LPS-primed BMDMs were transfected with flagellin (500 μg/mL) for 6 hours, intracellular taurine were detected (left). Immunoblots of active IL-1β (p17) and caspase-1 (p20) of these BMDMs in the presence of taurine (right)**.** Data are expressed as the mean ± SEM.

Fig. S4_Revision 1_02242025

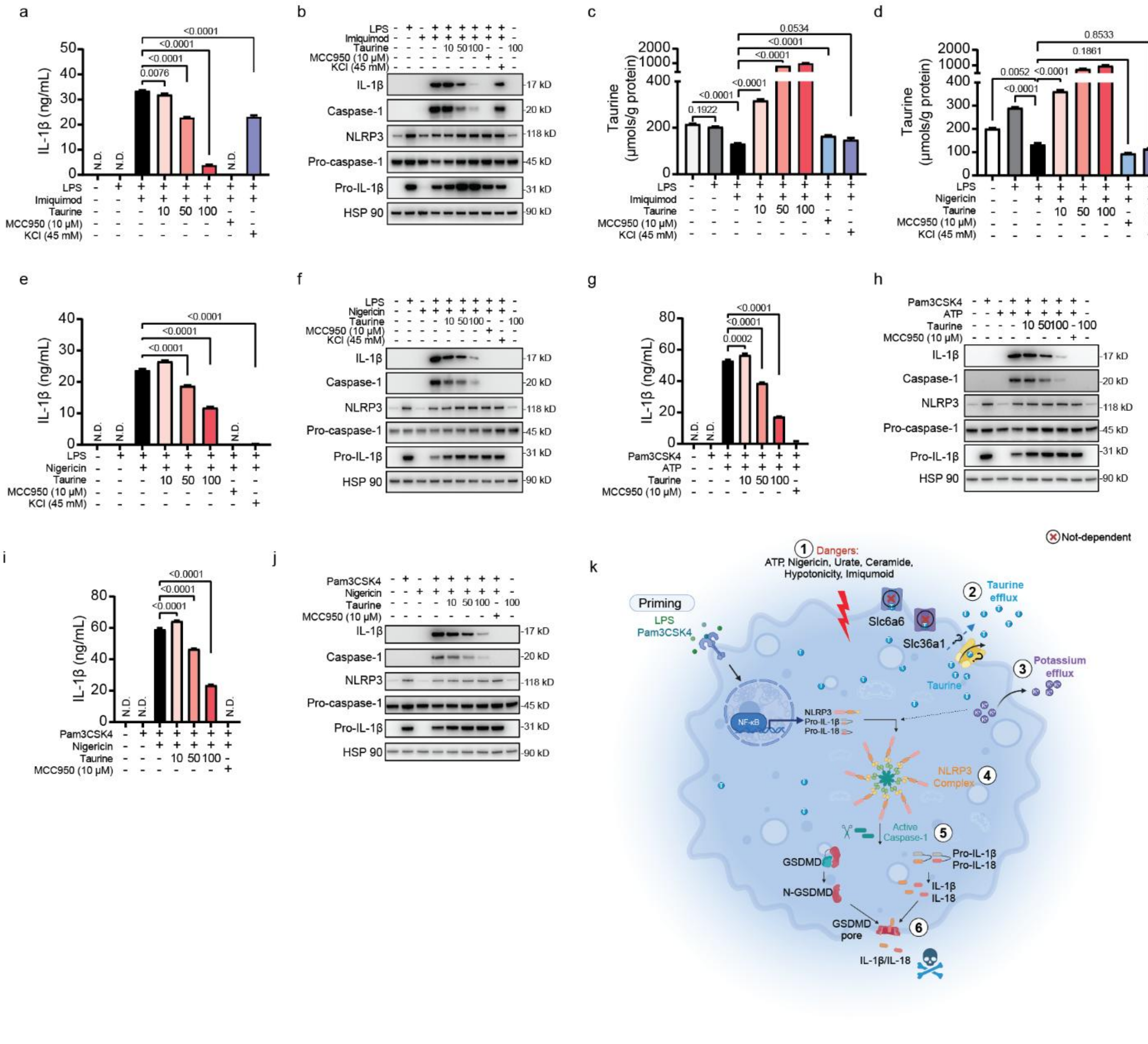


**Fig.S4. Taurine controls inflammasome activation upstream of NLRP3 and Potassium efflux.**

**(a-c)** BMDMs were primed by LPS for 4 hours followed by imiquimod treatment for 2 hours in the presence of taurine, MCC950 or KCl**.** (a) Production of IL-1β measured by ELISA. (b)

Immunoblot of caspase-1 (p20), IL-1β (p17) and NLRP3 in cell lysates of BMDMs. (c) Intracellular taurine levels in these BMDMs. These results are representative of three independent experiments. **(d-f)** BMDMs were primed by LPS for 4 hours followed by nigericin treatment for 1 hour in the presence of taurine, MCC950 or KCl**.** (d) Intracellular taurine levels in these BMDMs. (e) Production of IL-1β measured by ELISA. (f) Western blots of caspase-1 (p20) and IL-1β (p17) in BMDMs. These results are representative of three independent experiments. **(g-h)** BMDMs were primed by Pam3CSK4 for 4 hours followed by ATP treatment for 45 mins in the presence of taurine or MCC950**.** (g) Production of IL-1β measured by ELISA. (h) Western blots of caspase-1 (p20) and IL-1β (p17) in BMDM. (**i-j**) BMDMs are primed by Pam3CSK4 for 4 hours followed by nigericin treatment for 1 hour in the presence of taurine or MCC950**.** (i) Production of IL-1β measured by ELISA. (j) Western blots of caspase-1 (p20) and IL-1β (p17) in BMDMs. **(k)** A graphic abstract of the role of taurine in NLRP3 inflammasome activation. LPS- or Pam3CSK4-primed BMDMs generate the substrates for the NLRP3 inflammasome, including NLRP3, pro-IL-1β, and pro-IL-18. Various secondary signals induce taurine efflux followed by potassium ($K^+$) efflux, which triggers NLRP3 inflammasome assembly and activation of caspase-1. Active caspase-1 cleaves F-GSDMD, pro-IL-1β, and pro-IL-18. The cleaved N-GSDMD forms pores in the membrane, through which the active IL-1β and IL-18 are released. Canonical taurine transporters (Slc6a6 and Slc36a1) in macrophages are not required for taurine's inhibitory effects on inflammasome-dependent IL-1β.

Fig. S5_Revision 1_02242025

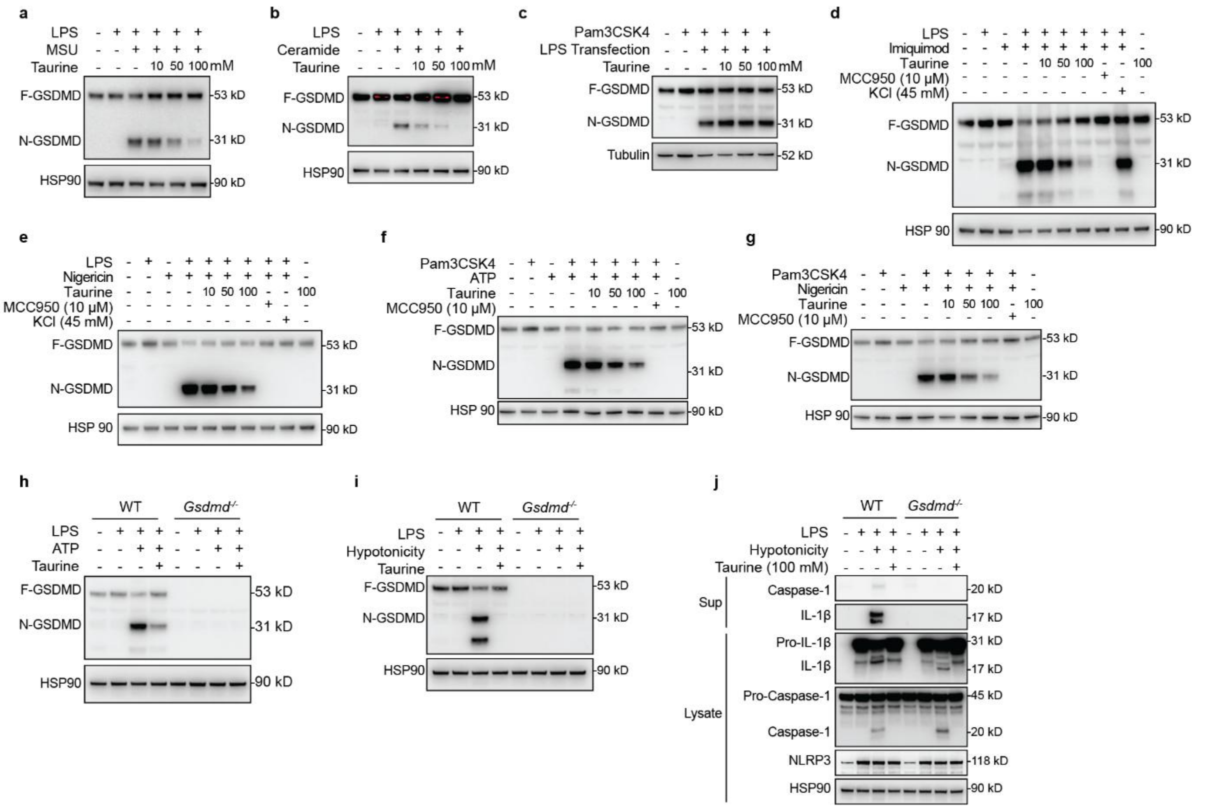

**Fig. S5. Taurine blocks Gasdermin-D (GSDMD) cleavage in response to NLRP3 inflammasome activation.**

**(a-b)** Western blots of GSDMD cleavage in cell lysates from BMDMs stimulated with LPS and MSU (a), Ceramide 6 (b) in the presence of taurine. These results are representative of three independent experiments. F-GSDMD, Full length GSDMD; N-GSDMD, N-terminal GSDMD. **(c)** Western blots of GSDMD cleavage in cell lysates from BMDMs stimulated with Pam3CSK4 and transfected with LPS in the presence of taurine. These results are representative of three independent experiments. **(d-e)** Western blots of cell lysate from BMDMs treated with LPS and imiquimod (d) or nigericin (e) in the presence of taurine, MCC950 or KCl. **(f-g)** Western blots of cell lysate from BMDMs treated with Pam3CSK4 and ATP (f) or nigericin (g) in the presence of taurine or MCC950. **(h-i)** Western blots of cell lysates from LPS-primed *Gsdmd*$^{-/-}$ BMDMs stimulated with ATP (h) or hypotonic solution (i) in the presence of taurine. These results are representative of three independent experiments. **(j)** Western blots of cell lysates and supernatants from *Gsdmd*$^{-/-}$ BMDMs stimulated with LPS and hypotonic solution and with taurine. These results are representative of three independent experiments.

Fig. S6_Revision 1

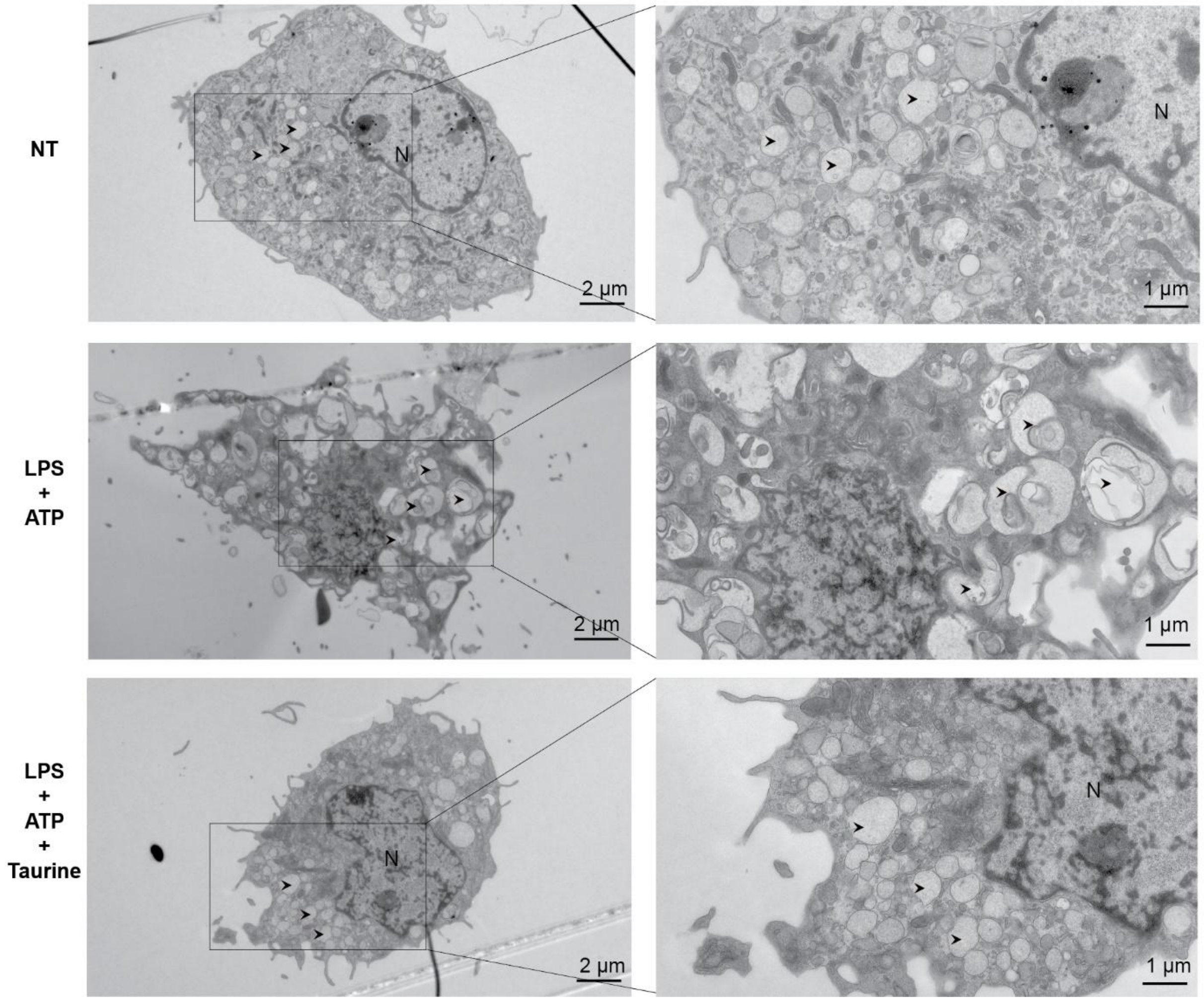

**Fig. S6. Taurine protects against pyroptosis.**
Resting macrophages containing autophagosomes with clear membrane structure (Upper). In response to LPS and ATP stimulation, macrophages containing autophagosomes with apparent remnants of degenerating parts (middle). With taurine addition, autophagosome with clear autophagosome vacuoles (lower). Scale bar: 2 µm in lower magnification (1,900 ×); 1 µm in higher magnification (4,800 ×). Arrowheads point to autophagosome. N, nucleus.

Figure S7_Revision 1

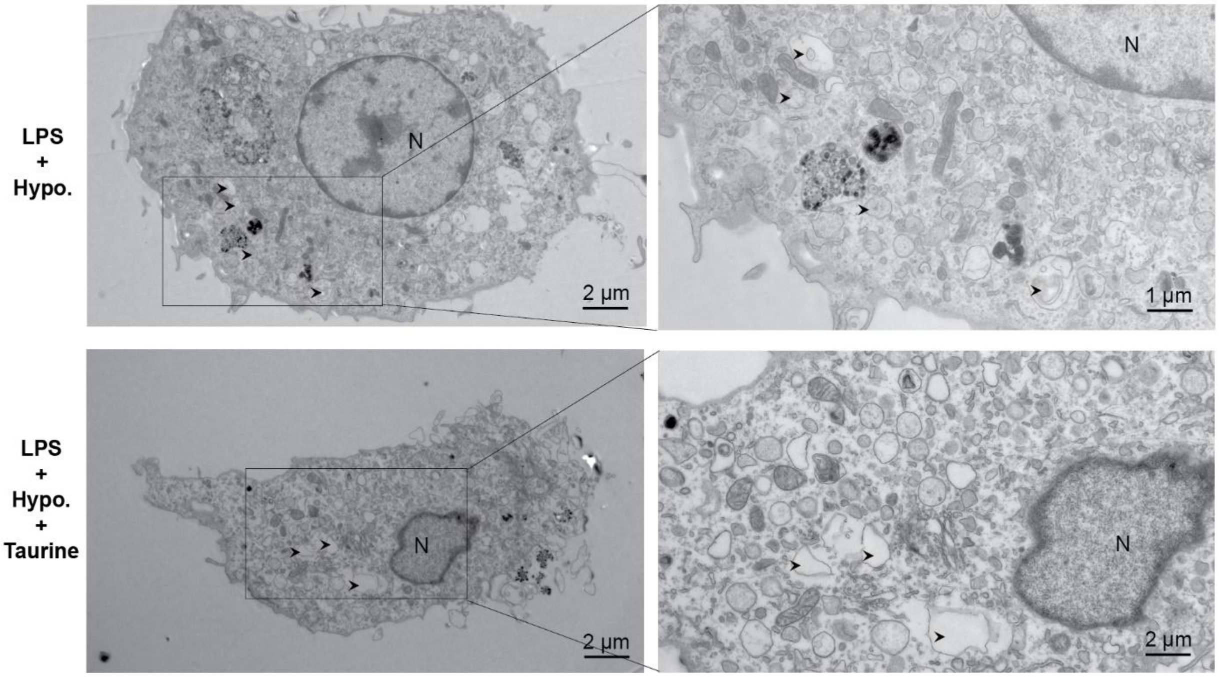


**Fig. S7. Hypotonicity induced cell swelling is reduced in the presence of taurine.**
Hypotonic solution induced cell swelling in LPS-primed BMDMs (Upper). With taurine supplement BMDMs decreased size (lower). Scale bar: 2 µm in lower magnification (1,900 ×); 1 µm in higher magnification (4,800 ×). Arrows point to the autophagosome. Hypo., Hypotonic solution, N, nucleus.

Fig. S8_Revision 1_02242025

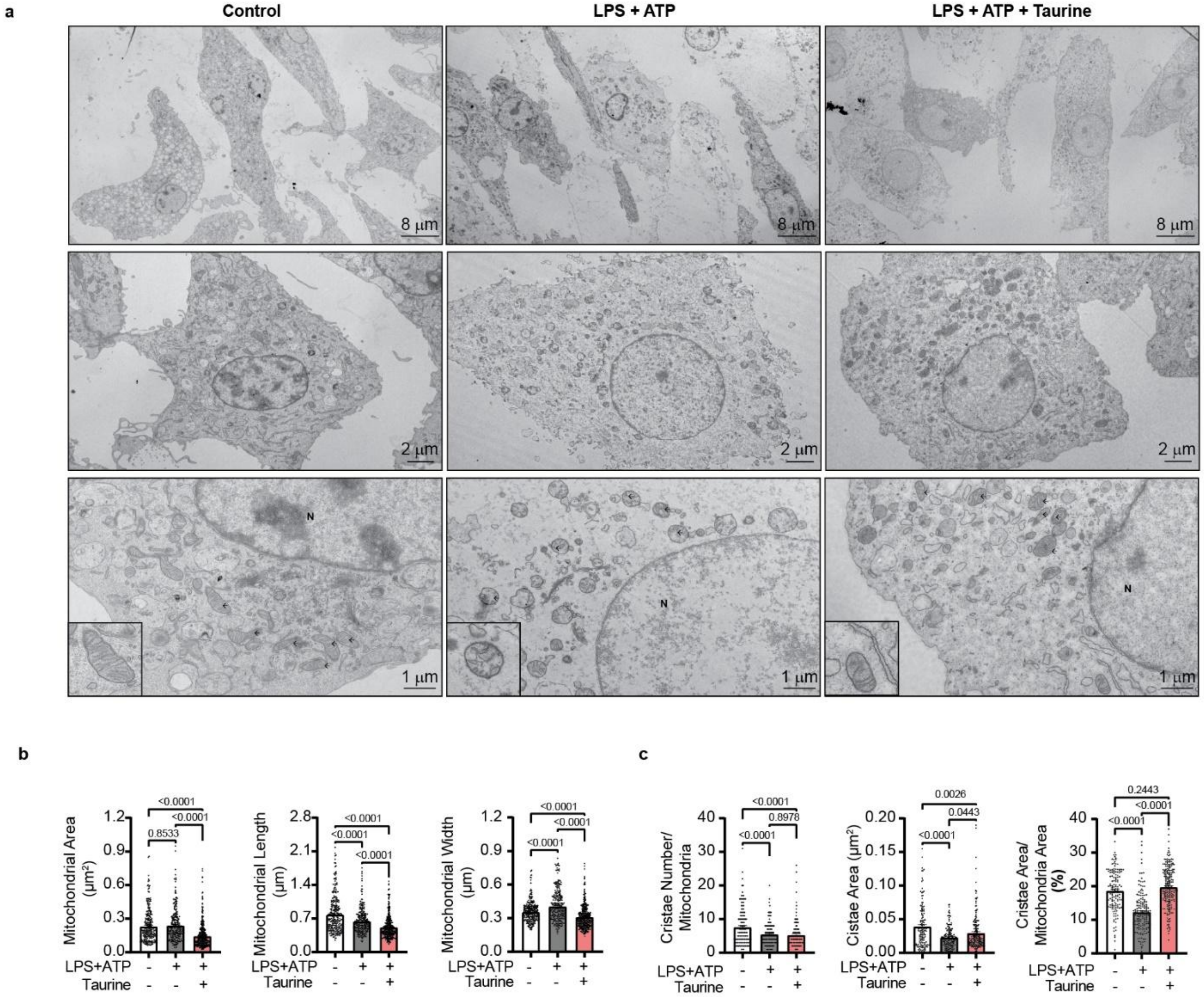


**Fig. S8. Taurine protects against inflammasome induced changes in mitochondria.**
**(a)** Representative TEM images of BMDMs after LPS (4 h) and ATP (30 min) treatment in presence of taurine. The lower magnification micrographs showed the status of cell population (Upper, scale bar: 8 μm, 690x). An individual BMDM was showed in the middle (scale bar: 2 μm, 1,900x). The lower panel displays the mitochondrion within the cytoplasm, with a representative mitochondrion illustrated in the left corner (scale bar: 1 μm, 4, 800x). Black arrow: mitochondrion; N: Nucleus. **(b)** Quantification of mitochondrial area, mitochondria length and width. **(c)** Quantification of cristae number, cristae area and cristae area/mitochondrial area. Error bars represent the mean ± SEM.

Fig. S9_Revision 1_02242025

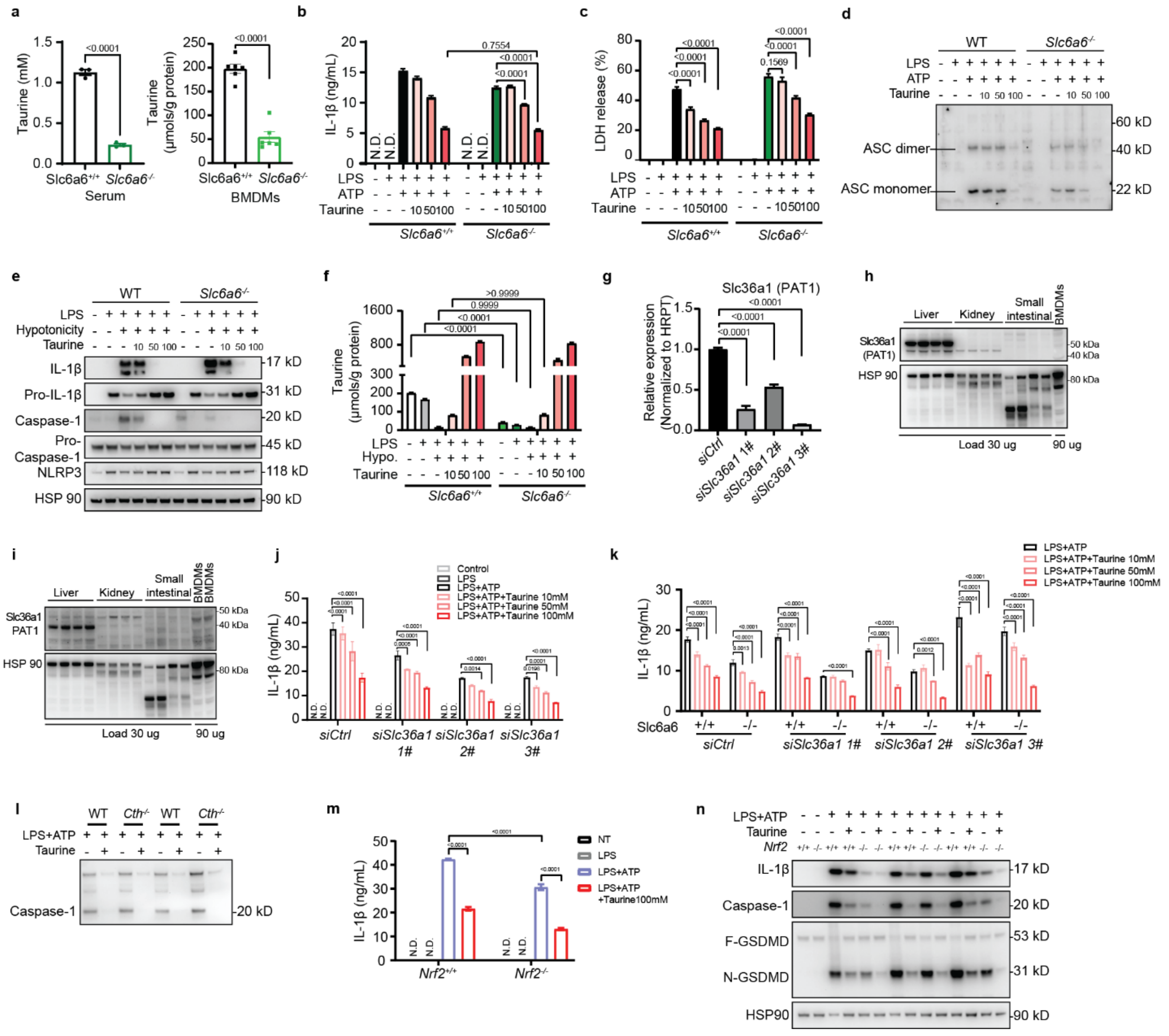


**Fig. S9. Taurine inhibits NLRP3 inflammasome independently of Slc6a6 or Slc36a1 transporter.**

**(a)** Taurine concentrations in the serum (n=3) and BMDMs (n=6) of control and *Slc6a6*$^{-/-}$ mice were determined. **(b-c)** Production of IL-1β (b) and LDH release (c) from LPS-primed *Slc6a6*$^{-/-}$ BMDMs stimulated with ATP and taurine. **(d)** Immunoblot analysis of ASC in cross-linked cytosolic insoluble pallets from *Slc6a6*$^{-/-}$ BMDMs stimulated with LPS and ATP in the presence of taurine. **(e)** Western blots of IL-1β (p17) and caspase-1 (p20) in cell lysates of control and *Slc6a6*$^{-/-}$ BMDMs stimulated with LPS and hypotonic solution in presence of taurine. These results are representative of three independent experiments. **(f)** Intracellular taurine levels of LPS-primed *Slc6a6*$^{-/-}$ BMDMs stimulated with hypotonic solution in presence of taurine. **(g)** Q-PCR analysis of *Slc36a1* levels of BMDMs transfected with siRNA of Slc36a1. **(h-i)** Immunoblots of Slc36a1 in different tissues and BMDMs using two different commercial antibodies. **(j)** Production of IL-

1β from *Slc6a6*$^{-/-}$ BMDMs transfected with siRNA for *Slc36a1*. LPS-primed cells were stimulated with ATP and taurine. **(k)** IL-1β production from wild-type and *Slc6a6*$^{-/-}$ BMDMs with RNAi knockdown of Slc36a1 and treated with LPS and ATP with or without taurine. **(l)** Western blots of caspase-1 of concentrated supernatants from LPS-primed *Cth*$^{-/-}$ BMDM stimulated with ATP in the presence of taurine. **(m)** Production of IL-1β (as measured by ELISA) from *Nrf2*$^{-/-}$BMDMs stimulated with LPS and ATP and treated with taurine. **(n)** Western blots of cell lysates and supernatants from *Nrf2*$^{-/-}$ BMDMs stimulated with LPS and ATP and with taurine. Data are expressed as the mean ± SEM of three independent experiments carried out in triplicate. Error bars represent the mean ± SEM.

Fig. S10_Revision_02242025

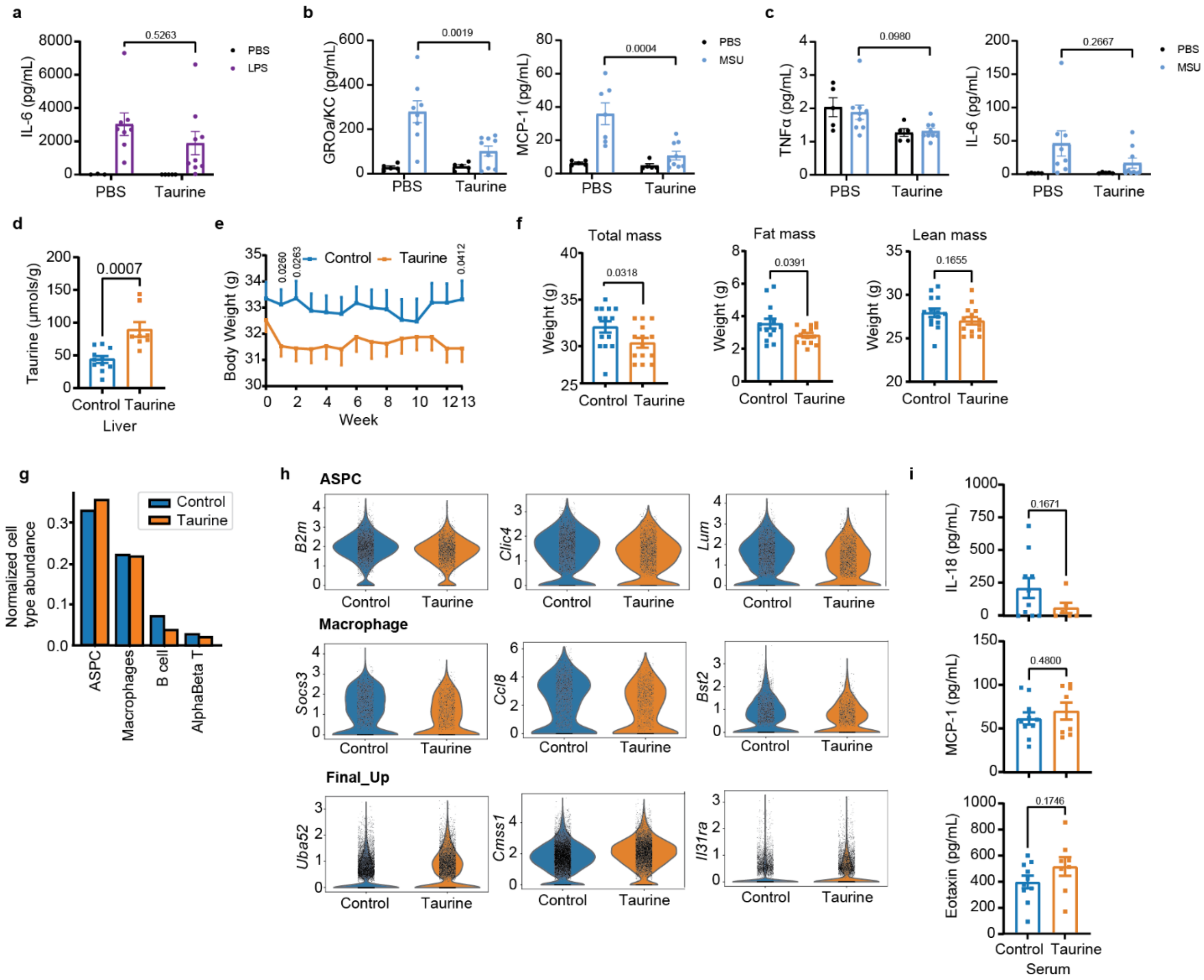


**Fig. S10. Taurine supplementation inhibits inflammation *in vivo*.**

**(a)** Serum levels of IL-6 from 8- to 9-week-old C57BL/6J male mice pretreated with taurine or vehicle control as measured by ELISA 4 h after i.p. LPS injection. (n=3, 8, 4, 9). **(b)** Serum levels of GROα and MCP-1 from 8-week-old C57BL/6J male mice pretreated with taurine or vehicle control as measured by ELISA 6 h after i.p. MSU injection. (n=5, 8, 5, 9). **(c)** Serum levels of TNFα and IL-6 from 8-week-old C57BL/6J male mice pretreated with taurine or vehicle control as measured by ELISA 6 h after i.p. MSU injection. (n=5, 8, 5, 9). **(d-i)** Aged male C57BL/6N mice (20-month-old) given normal water or drinking water with taurine (8,000 mg/kg/day, 4% in water (w/v)) for 4 months. (d) Taurine levels in liver after 4 months of taurine supplementation. (e) Body weight curves of aged mice with or without taurine supplementation for 13 weeks. (f) Total mass, fat mass and lean mass measured by EchoMRI of these male C57BL/6N mice after 3 months of taurine supplementation (n=14,14). (g) Main cell composition in single cell RNA-sequencing of mice VAT after 4 months of taurine supplementation. (h) Significantly down-regulated genes in adipocyte stem and progenitor cells (ASPCs), and macrophages by scRNA-sequencing (top and middle panel). Significantly up-regulated genes in all cells in VAT (lower panel). (i) Serum levels of IL-18, MCP-1 and Eotaxin from these aged male C57BL/6N mice treated with taurine for 4 months (n=9, 8). Error bars represent the mean ± SEM.
